# Neural dynamics indicate parallel integration of environmental and self-motion information by place and grid cells

**DOI:** 10.1101/640144

**Authors:** Dmitri Laptev, Neil Burgess

## Abstract

Place cells and grid cells in the hippocampal formation are thought to integrate sensory and self-motion information into a representation of estimated spatial location, but the precise mechanism is unknown. We simulated a parallel attractor system in which place cells form an attractor network driven by environmental inputs and grid cells form an attractor network performing path integration driven by self-motion, with inter-connections between them allowing both types of input to influence firing in both ensembles. We show that such a system is needed to explain the spatial patterns and temporal dynamics of place cell firing when rats run on a linear track in which the familiar correspondence between environmental and self-motion inputs is changed (Gothard et al., 1996b; Redish et al., 2000). In contrast, the alternative architecture of a single recurrent network of place cells (performing path integration and receiving environmental inputs) cannot reproduce the place cell firing dynamics. These results support the hypothesis that grid and place cells provide two different but complementary attractor representations (based on self-motion and environmental sensory inputs respectively). Our results also indicate the specific neural mechanism and main predictors of hippocampal map realignment and make predictions for future studies.

## 1. Introduction

The place cells in the rat hippocampus show strong behavioral correlates by firing only when the animal visits a particular localised region of the surrounding environment (O’Keefe and Dostrovsky, 1971; O’Keefe, 1976). Collectively these place cells provide a population code for spatial position. A neural representation (‘cognitive map’) of a particular environment is formed in such a way (O’Keefe and Nadel, 1978). As the animal moves around a particular environment, the firing pattern of place cells is continuously updated, reflecting the current position of the animal. This continuous shifting of neural representation could be driven by at least two types of information - perceptual from the environment and internally generated concerning the rat’s own movements, and takes place even in total darkness (O’Keefe, 2007).

Another type of spatially selective cells was found in the subiculum by Lever et al. (2009), and is referred to as Boundary Vector Cells (BVCs), due to the fact that a particular BVC fires maximally when a boundary is encountered at the BVC’s preferred distance and allocentric direction from the rat. ‘Border cells’, fulfilling the criteria for BVCs, have also been found in the mEC and adjacent parasubiculum by Solstad et al. (2008), and may represent a subset of BVCs. Initially, the model that incorporated putative BVCs as providing sensory inputs to place cells was developed by Hartley et al. (2000), based on the findings from earlier studies of O’Keefe and Burgess (1996). Barry et al. (2006) then further demonstrated experimentally that impediments to movement, whether walls, a free standing barrier or a sheer drop, play a key role in defining place cell firing, and predicted existence of cells in the subiculum that fit important elements of the BVCs in the Hartley et al. (2000) model. Recently, Grieves et al. (2018) used a similar BVC model of place cell firing to replicate place field repetition seen in various experiments with multicompartment environments.

Updating of place cells’ activity according to the rat’s internally generated motion signals could be achieved via grid cells, another type of spatially selective cell, that fire whenever a rat enters one of an array of locations arranged in a hexagonal grid across the environment (Hafting et al., 2005) and provide anatomical inputs to areas CA3 and CA1 of the hippocampus. Since grid cells preserve the shape and size of their grid-like firing patterns despite removal of visual cues, it is possible that a path integration mechanism is responsible for maintenance of the grid structure. The fact that neighbouring grid cells have the same field size and spacing, as well as slightly offset grid phasing (Hafting et al, 2005), and that recurrent connections are present in the mEC (Lingenhohl and Finch, 1991; Germroth et al, 1991; Dhillon and Jones, 2000) suggest that grid cells may perform path integration via a continuous attractor based mechanism (McNaughton et al., 2006). At the same time, a number of experiments have shown an influence of environmental boundaries also on grid cell firing. For example, in the experiments by Barry et al. (2007) and Stensola et al. (2012), the environment deformation caused partial rescaling of grid cell firing patterns, while the experiments by Krupic et al. (2015) and Stensola et al. (2015) showed effects of alignment between grid patterns and boundaries. Recently, Krupic et al. (2018) also found that local changes to the configuration of the enclosure lead to localized changes in the grid structure (together with local changes in place and boundary cells’ responses).

In this study we investigate whether the integration of environmental and self-motion information into a representation of an estimated position could result from a reciprocal interaction between recurrent networks of place cells (receiving inputs from BVCs) and grid cells (implementing path integration) (O’Keefe and Burgess, 2005). We test our hypothesised model against experimental data on place cell firing in situations where sensory and self-motion information are put into conflict (Gothard et al., 1996b; Redish et al., 2000). The Gothard et al. experiment was previously simulated by Sheynikhovich et al. (2009) using a model that integrates visual and self-motion information in its grid cell population, and recently by Keinath et al. (2018) using a model in which path integrating grid cells receive direct input from border cells. Importantly, however, these single-layer attractor models could not capture the neural dynamics observed in the experiment, as we discuss below.

Together with testing the hypothesis about the specific neural implementation of self-motion and sensory information integration, we also address the questions (initially raised by Redish et al., 2000) regarding the specific mechanisms and the main predictors of the hippocampal map realignment process. Finally, we compare the model to an alternative, single-layer continuous attractor model, and discuss its relationship to other related models in terms of their ability to explain the temporal dynamics of place cell firing in these experiments.

## 2. The Model

In our firing rate model, integration of self-motion occurs between grid cells and projections from grid cells to place cells provide the self-motion contribution to place cell firing. The sensory contribution to place cell firing comes through BVCs, and the projections from place cells to grid cells maintain the stability of grid cell firing relative to the environment. The animal’s location is hypothesised to be determined on the basis of these interactions.

### 2.1 Place cells with Boundary Vector Cell (BVC) environmental inputs

Place cells in area CA3 of the rat hippocampus receive inputs from the mEC, which contains grid cells, BVC-like border cells and itself receives inputs from the BVC-containing subiculum. CA3 in turn projects to CA1, another hippocampal area with a large place cell population. We assume that the place cells in CA1 represent direct feed-forward readout of CA3 place cells’ activity, and thus could be omitted from our model without significantly affecting its overall dynamics.

A characteristic anatomical feature of the CA3 region of the hippocampus is the extensive recurrent connections between its pyramidal cells (Amaral & Witter, 1989). The presence of recurrent connections suggests that the network may be subject to stable attractor dynamics, which means that place cell activity patterns correspond to the stable equilibria states of a potential CA3 attractor network. A number of researchers to date have taken an attractor dynamics approach to modelling the behaviour of recurrent networks (Zhang, 1996; Samsonovich & McNaughton, 1997; Redish & Touretzky, 1998; Stringer et al, 2002b; Conklin & Eliasmith, 2005), referred to as ‘continuous attractor’ networks, since they can stably maintain patterns of firing of their neurons corresponding to any location in a continuous physical space, forming a whole Cartesian plane of stable fixed points (a ‘plane attractor’).

In our model, we use a 2-D sheet of recurrently connected place cells, arranged so that each one’s location in the sheet (covering the simulated full track environment) corresponds to its preferred firing location in the environment, so that the firing pattern over the neural ensemble forms an ‘activity bump’. The connection strength between any two neurons is inversely related to the distance between their locations, which, together with a global feedback inhibition, produces a plane attractor (see the Appendix for details).

Every place cell in the model is assumed to receive inputs from four orthogonally tuned BVCs, each of which has directional tuning perpendicular to one of the four surrounding boundaries. Our estimated BVC tuning curves (see *‘BVC inputs to place cells’* in the Appendix for details) differ from the Gaussian tuning curves used by O’Keefe and Burgess (1996) to model inputs to individual place cells in that the standard deviation of a particular (otherwise Gaussian) curve is not constant, but instead is a linear function of the distance from the peak of the curve. See Figure 1. This makes our curves skewed, so that the side oriented towards the boundary of interest is steeper than the one oriented away. Such a shape seems to be physiologically plausible, since shorter distances are easier to estimate.

**Figure 1.**
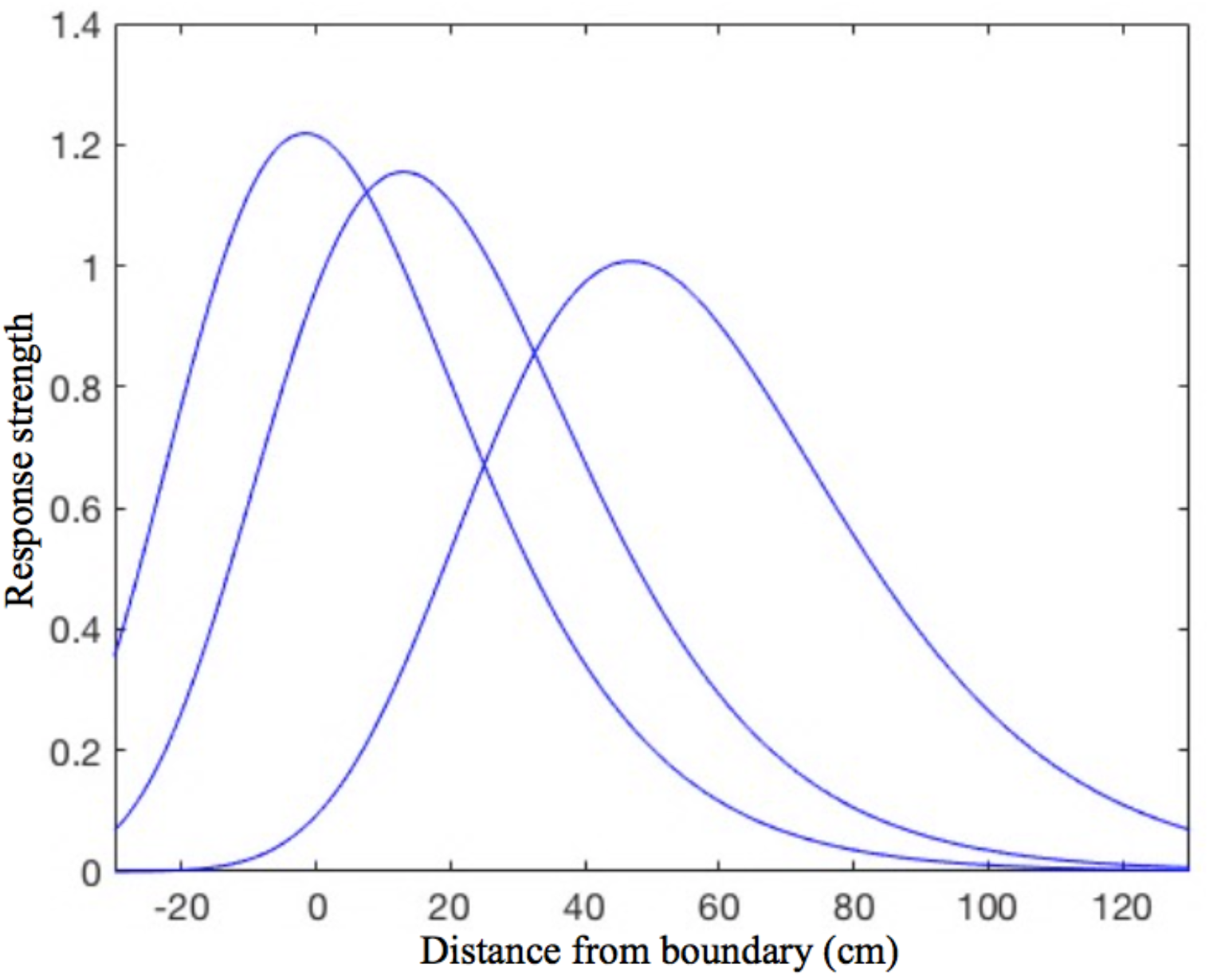
The shape of the BVC tuning curves in our model. Note the slight asymmetry, as the curves standard deviation is not constant, but increases linearly with increasing distance of the rat from the boundary. The distance tuning is narrower for cells which have a peak response near to boundaries and gradually widens with the distance of peak response. This is consistent with the rat being able to judge shorter distances more accurately, which also implies that the BVCs tuned to shorter distances exert more influence on place cell firing (which is reflected in the curves height).

### 2.2 Grid cells and path integration

Each grid cell fires in multiple locations, arranged as the nodes of an equilateral triangular grid spanning the whole environment (Hafting et al., 2005). The field sizes and spacing of the grid cells located more dorsally/ventrally in the mEC differ, increasing with depth from the postrhinal border in a stepwise manner indicating the presence of discrete modules (Barry et al., 2007; Stensola et al., 2012). Neighbouring grid cells have the same field size, orientation and period of grids, but their grids are offset in phase. This indicates that conjunctions of active neurons are repeated periodically as the rat moves over a surface, so that the whole environment is covered by the nodes of a local cell module with a common grid spacing and orientation. This, together with the presence of recurrent connections in the mEC (Lingenhohl and Finch, 1991; Germroth et al., 1991; Dhillon and Jones, 2000), suggests that modules of cyclically organized grid cells may perform path integration via a continuous attractor based mechanism (McNaughton et al., 2006; Guanella et al., 2007; Burak and Fiete, 2009).

In order for the continuous attractor neural network to perform path integration requires the presence of asymmetric synaptic pathways between grid cells with firing patterns offset in 6 directions (defined by the axes of the cells hexagonal firing pattern). These pathways need to transmit activity determined by the agent’s movement direction and speed, as suggested for place cells (and for which fewer than 6 directions can be used, Zhang, 1996; Samsonovich and McNaughton, 1997; Conklin and Eliasmith, 2005). We assume such pathways are mediated by the conjunctive grid by head-direction cells that have been found in the mEC by Sargolini et al. (2006) (see Equations (A.9) and (A.10)), whose output we assume to also be modulated by running speed.

Grid cells are thought to project to place cells, which provide the means of combining inputs from grid cell modules with different scales into a unified path integration–based estimate of position. A particular place cell receives connections from all the grid cells, with various grid spacing, that happen to be active near to the place field in a particular environment. The connections strength is inversely related to the distance between the place and grid cells preferred firing locations, similarly to the recurrent connections among the place or grid cells. The combined input from all the grid cells connected to a particular place cell will be maximal at the centre of the place field and will decay with increasing distance from it, since the inputs from cells with different grid scales will no longer converge. Combining the output of grid cells from multiple modules also allows a significantly larger representational capacity than a single network with the same overall number of cells would allow (Fiete et al., 2008).

Continuous attractor models of grid cell networks have been shown to be capable of accurate path integration over comparable distance and time-scales to those probed in behavioral assays (Burak and Fiete, 2009), and are preferable to single cell models since they are more robust to the presence of noise (Navratilova et al., 2011). Yet even noise-free, large networks have only finite integration accuracy (Burak and Fiete, 2009), and potential errors in their velocity input would also contribute to an increasingly incorrect position estimate over time. Thus, perceptual environmental information is required to maintain the grid cell-based representation of location in register with the environment. In the absence of such information, behavioural measures of path integration demonstrate rapid increase in error with movement (Etienne, Maurer and Seguinot, 1996). In order to anchor the grid cell firing patterns to the environment, place cells are presumed to project to grid cells, with the connections strength being inversely related to the distance between their preferred firing locations. This anchoring cannot be achieved so easily by providing sensory inputs, e.g. from the BVCs, directly to grid cells, since their responses are not constrained to one specific location in the environment. The same place cell – grid cell connections also serve the purpose of registering together different modules of grid cells. This is needed in order to maintain a stable relationship between them, and cannot be achieved by connecting these modules directly since the firing of two grid cells from different modules might only overlap at a single location in an environment.

### 2.3 The place cell – grid cell model

In order to model the behavior of the navigation system based on the reciprocal interactions between place cells and grid cells, we assume (for simplicity) that inputs to the layer of place cells are provided by three modules of grid and grid by head-direction cells, each with different grid scaling. Each module includes 441 grid cells, all recurrently connected in a manner that could result from Hebbian learning from the hexagonal topology of multiple firing fields of individual cells. The grid scale of each successive module is a factor 0.72 smaller (or 1.39 larger) than the next, which is similar to the ratio found by Stensola et al. (2012). The standard deviations of the individual firing fields scale accordingly, consistent with physiological data, as follows: 5cm, 7cm and 9.65cm. The standard deviation of the place cell firing fields is 7cm. The strength of synaptic connections from both the grid cells to place cells, and the place cells to grid cells, is defined by a Gaussian function of the distance between the grid and place cell preferred locations, with the variance of the Gaussian given by the average of the variances of the place and grid cell firing fields. The outlined system is represented by the set of equations (A.11) in the Appendix.

## 3. Materials and Methods

### 3.1 Neural population dynamics, Gothard et al. (1996b) and Redish et al. (2000)

Gothard et al. (1996b) recorded populations of hippocampal neurons in rats shuttling on a linear track between a movable reward site, mounted in a sliding start box, at one end and a fixed reward site at the opposite end. The rats were pretrained on the full-length track with a constant box position. On each subsequent trial, the box was randomly moved to one of five equally spaced locations, thereby creating mismatches with the originally learned relationships of the box to other cues in the environment (Figure 2). The movement of the box took place while the rat ran ‘outbound’ toward the fixed reward site, with the rat then returning ‘inbound’ to the box in its new position. Along a journey, the same cells were active, in the same order, regardless of the box location, although elements of the sequence of place fields on the full track were sometimes omitted.

**Figure 2.**
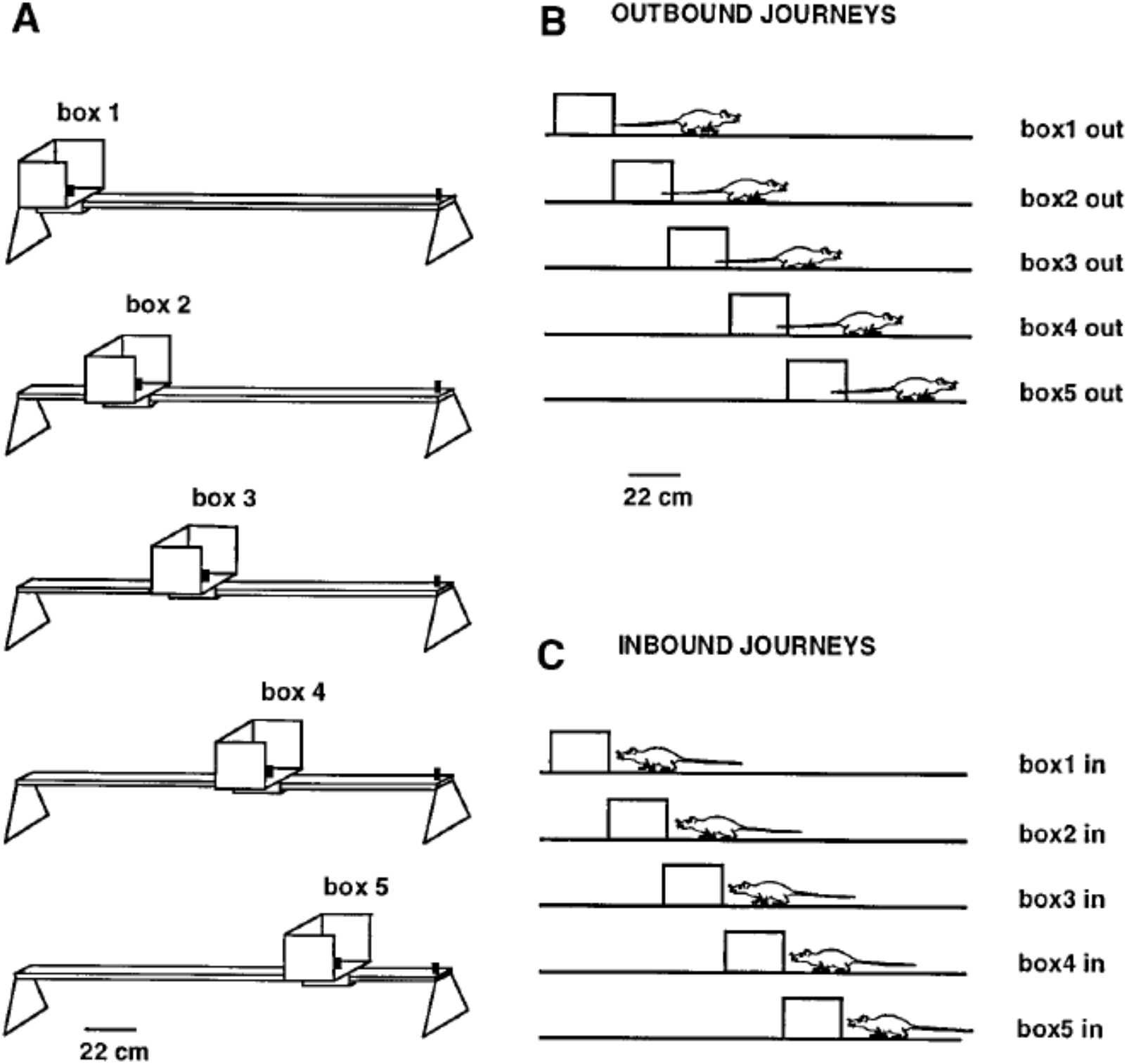
*A*, Linear track (161cm long outside the movable box) with the five equally spaced box locations used as the start or end point of each journey. *B* shows the five types of outbound journeys, labeled *box1 out*, *box2 out*, etc. *C* shows the five types of inbound journeys. Adapted from Gothard et al. (1996b).

The rat’s internal spatial representation, defined by place cell population activity, was quantified in terms of population vectors. Then the similarity of the population activity on the full-length journey to the population activity on each of the four types of shortened journeys was tested by correlating point by point the population vectors computed for each spatial location. The results of these correlation procedures for a single rat are represented graphically in Figure 3, adapted from Gothard et al. (1996b).

**Figure 3.**
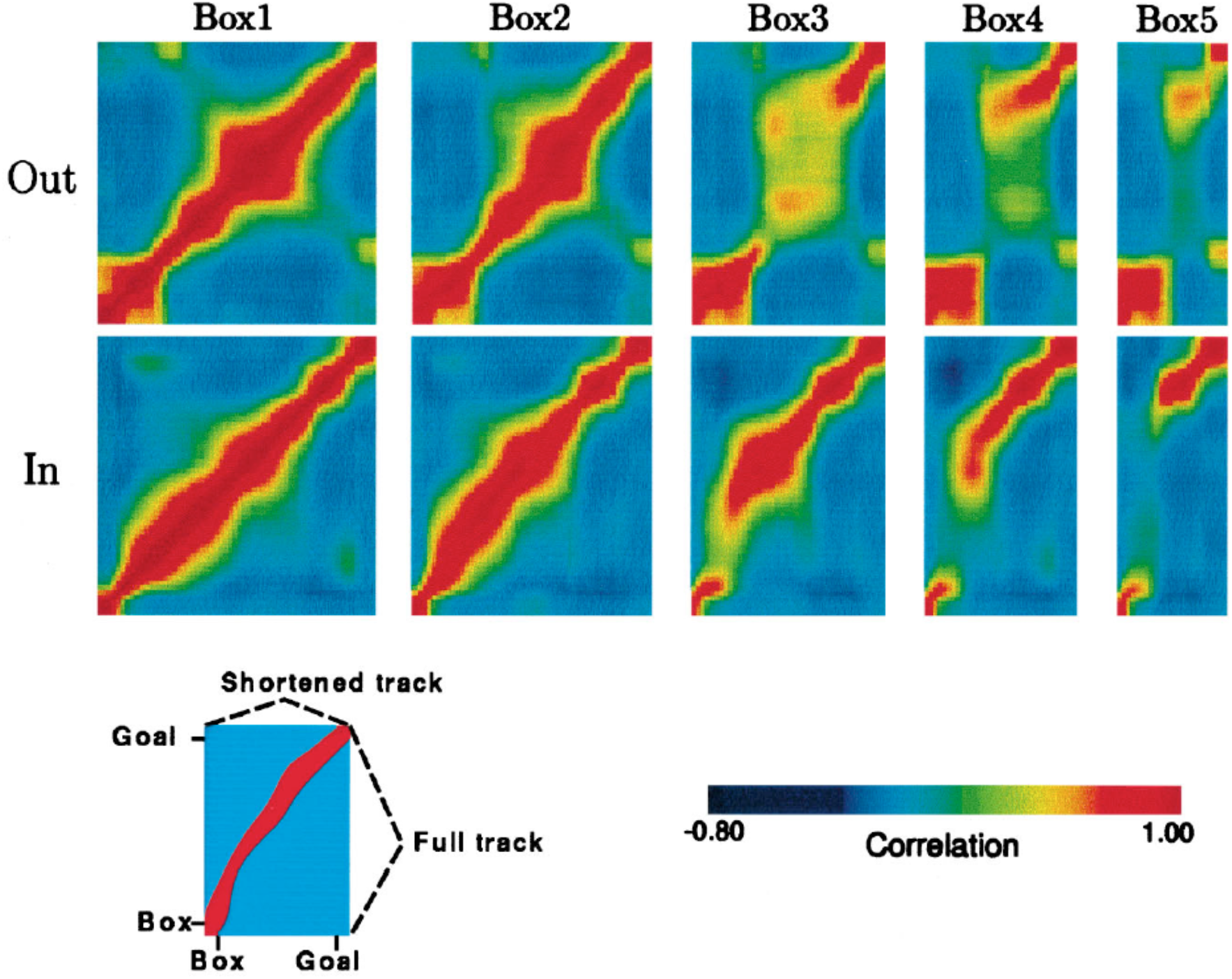
Population vector correlations between the pattern of firing on the full track and on the shortened tracks for one rat, whose pattern is representative of all other rats except one. Population vector correlations are shown for the outbound journeys (top five plots) and inbound journeys (bottom five plots). For each correlation plot, the vertical axis corresponds to the full track (including the box), whereas the horizontal axis corresponds to the length of the track (including the box) covered by the rat in one of the five trial types, i.e. box1, box2, etc (see key, bottom left). Highly correlated firing patterns between one location on the full track and a second location on one of the shortened tracks are indicated in *red*. The first plot in each row is a spatial autocorrelation of the population vectors on the full track, with values of 1.0 along the diagonal. For the journey types 4 and 5, when the track was greatly shortened, the discontinuity of the ridges can be seen, which is due to the ‘jumping’ of the activity packet over the intermediate stages.

The principal finding was that when a mismatch existed between the rat’s internal spatial representation and the rat’s coordinates in the external reference frame centered on the reward site the rat was running to, a dynamic correction process took place. The correction always took place after some initial delay, which was longer for the longer outbound journeys and shorter for the shorter ones. On longer tracks with moderate mismatches the activity realignment usually occurred already past the midpoint, in the second half of the journey. On some outbound journeys the internal representation remained more aligned with the distance from the box for about 1 meter, even though the start box was behind the rat, outside its field of view. For moderate mismatches, the internal representation, after some initial delay, was shifted continuously through intervening states, faster than the actual speed of rat, until it was closely aligned with the landmarks corresponding to the end of the track the rat was approaching. The shorter the track, the more rapid the correcting shifting of the activity. In case of large mismatches, however, the internal representation jumped abruptly to the new position, skipping the intervening states, which occurred in the first half of journey, before the midpoint between the front of the start box and the end boundary.

Based on the results of this work and also of their earlier study (Gothard et al., 1996a), in which rats shuttled between a box and a pair of landmarks placed variably in a large arena, Gothard et al. proposed that the firing of place cells is controlled by a competitive interaction between path integration and external sensory inputs, primarily vision.

In Gothard et al.’s (1996b) experiment, there is a considerable difference between the outbound and inbound journeys. In case of the outbound journey, a change in the box location results in a change of the rat’s starting position relative to the entire room. In case of the inbound journey, on the contrary, the rat’s position in the room frame does not change, only one of the track boundaries changes its location. And this difference between the two journey types manifests in Gothard et al.’s data, with the population activity realignment on the inbound journeys occurring much later than on the outbound, just before the rat enters the box.

According to the difference in the realignment dynamics of the population activity on the outbound vs inbound journeys, the room features have a strong influence on the place cell activity. This could be primarily due to the room walls, since the track was placed across the corner of the room. Thus, on the inbound journey, for a large part of the journey (e.g. till the vicinity of the box) the sensory inputs from ahead of the rat may be coming predominantly from the room wall, rather than the mismatched box that is much smaller, which may explain the observed dynamics (i.e. the realignment to the reference frame of the box occurring just before it on all the track lengths). Because of this ambiguity, we focus on the outbound journeys, in which path integration from the start box mis-matches with constant environmental sensory inputs from all around the rat, excepting only the small start box directly behind it. Any sensory input from the box is likely cancelled out by the conflicting sensory input from the wall behind the rat in the overall sensory input from behind the rat. Therefore in our outbound runs simulations we assume that there is no sensory influence on the rat from the movable box.

In a subsequent study, Redish et al. (2000) randomly varied the shortened track lengths (by varying a movable box position) between 150cm and 90cm (as measured from the front of the box to the end barrier) across different trials, thus sampling different track lengths from within the range during a 30min experiment. The full track length was similar to that used by Gothard et al., and each trial also consisted of outbound (towards the end barrier) and inbound (back to the box) journeys.

The study replicated and extended that of Gothard et al. (1996b) through studying the realignment dynamics by measuring properties of the place cell ensemble activity and the observation of the realignment on a moment-by-moment basis within individual journeys. The major conclusion of the study was that the realignment of the ensemble activity occurs after a temporal delay, suggesting that there is a stochastic switch occurring somewhere in the system. We provide more details of the study in the ‘*Detailed analysis of* Redish et al.’s (2000) *data’* part of ‘*Results’*, where we analyse Redish et al.’s data using our simulation results.

### 3.2 Simulation methods

We simulate the situation of competitive interaction between path integration and external sensory inputs, corresponding to outbound runs in Gothard et al. (1996b) and Redish et al. (2000), using the place cell – grid cell model, represented by Equations (A.11). We vary the maximum strength of synaptic connections from grid cells to place cells, given by *G* in the model, in the range between 1 and 3, in order to investigate its influence on the model. The strength of synaptic connections from place cells to grid cells (*P*) we keep constant and equal to 10. It is set higher than the combined strength of the connections to place cells from the three grid cell modules to compensate for the increased threshold for synaptic transmission from place to grid cells (see the Appendix).

As explained in the part 3.1, we assign a weighting factor of zero to the inputs from the BVCs tuned to the start box on outbound runs. Therefore the total external input *h*_*i*_ to the place cell *i*, given by (A.8) in our model (A.11), is comprised of an input from the BVC tuned to the boundaries ahead of the rat and inputs from two BVCs tuned to the boundaries in the lateral directions. The sum of the three BVC inputs is thresholded using a fixed threshold *T* that we set equal to the sum of the two laterally tuned BVCs inputs, which remain constant along the track since the rat runs parallel to the track side borders. Each individual BVC *i* input *b*_*i*_ is given by (A.7), with the following parameters: *A*_*0*_ = 4.5, *σ*_*0*_ = 21.5, *α* = 0.109, *β* = 0.016, *γ* = 0.101.

During simulations the starting position of the rat was changed in such a way as to represent the ranges of track lengths in both Gothard et al.’s (1996b) and Redish et al.’s (2000)) experiments. Similar to those experiments, the full (familiar to the rat) track length was 160cm (outside the moveable starting box), and its four equally spaced shortened versions were 140cm, 120cm, 100cm and 80cm in length. The direction of the rat’s movement was *θ* = 0 (i.e. from the box front along the track to its end) and the speed *V* = 15 cm/s across all trials.

For comparison, using the same settings, we also simulate a ‘place cell only’ model in which there are no grid cells and only a single continuous attractor network. As for the place cell – grid cell model simulation, we use a similar 2-D sheet of recurrently connected place cells (a plane attractor) that covers the full length of the familiar 160cm track. Path integration is performed via asymmetric connections between the place cells, which strength is modulated by the rat’s direction and velocity signals. At the same time sensory inputs to the place cells are provided by BVCs, in the same way as in the main model (A.11). Further details of the place cell only model (A.12) implementation are given in the Appendix.

## 4. Results

### 4.1 Realignment dynamics favour the place cell – grid cell model

The behaviour of the place cell - grid cell model (A.11), with *G* = 2.1 (i.e. the grid cell – place cell connection strength that we varied between 1 and 3), provides a good qualitative fit to the behaviour of place cells on outbound journeys in Gothard et al.’s (1996b) study, across all simulated track lengths.

During moderate self-motion and sensory information mismatches (as on the longer two of the shortened tracks), after a pronounced initial delay, the place cell activity bump was continuously shifted through intervening positions until its location was in agreement with the sensory inputs provided by the BVCs tuned to the approaching end of the track. The speed of transition depended on the mismatch size, with a larger mismatch resulting in a more rapid transition, following an initial delay (Figure 4, *top*). When the mismatch was large (as on the two shortest tracks), the activity bump dissolved in its initial location and instead emerged in a ‘correct’ one, in line with Gothard et al.’s findings (Figure 4, *bottom*). Such jump realignments occurred quicker, in the first half of the journey (from the front of the box to the track end), whereas the continuous shift realignments occurred much farther from the start, in the second half. The shorter the track, and thus the nearer the start box to its end, the sooner the realignment occurred across all the track lengths.

**Figure 4.**
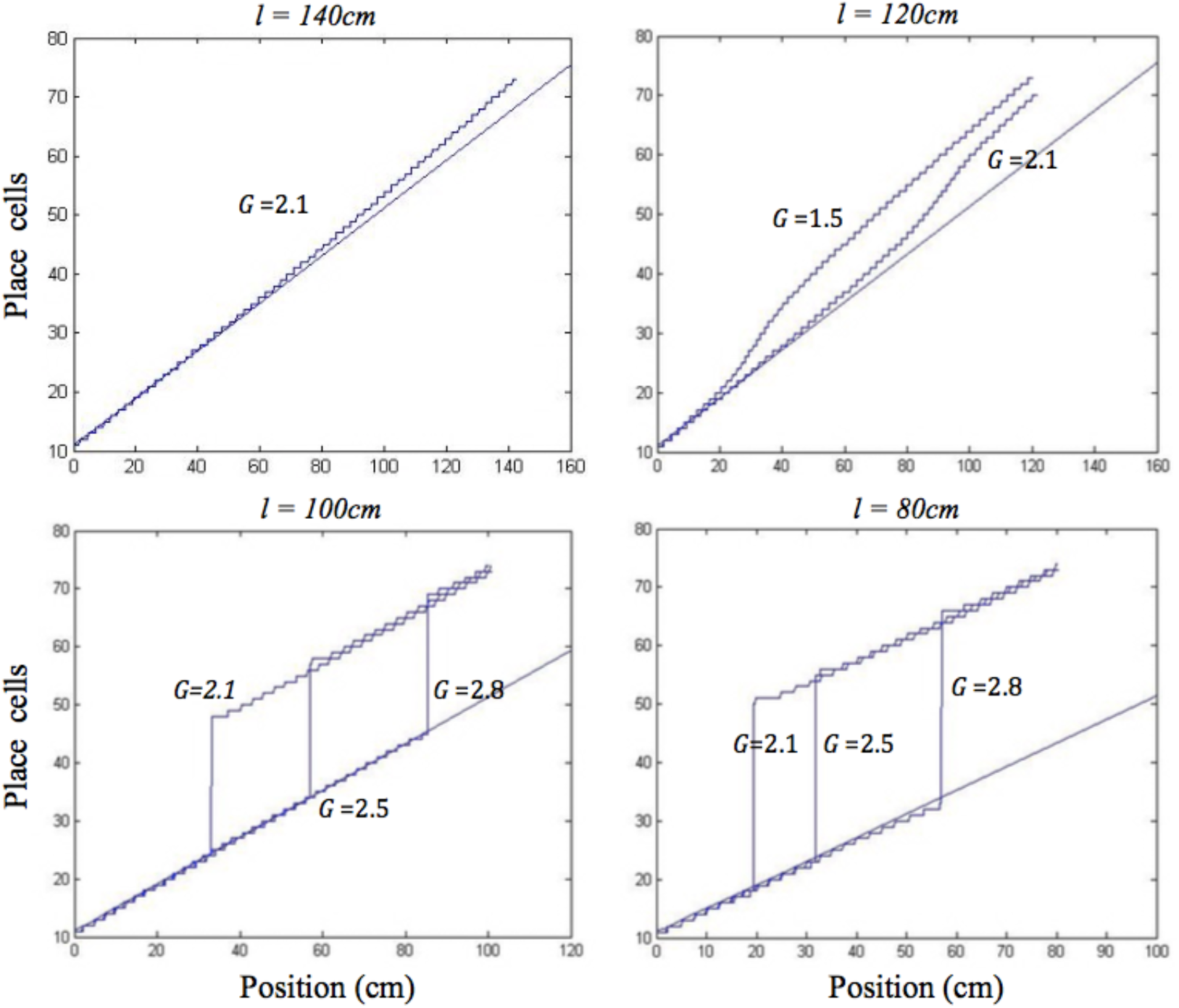
Realignment of the simulated place cell representation of location as track length varies in the place cell-grid cell model. Plots show position on the track on the *x* axis and the relevant place cells ordered by their location of peak firing on the full length track (160cm) on the *y* axis. The place cells in rows 11-75 have firing peaks evenly distributed along the full length track from the front of the box. The straight blue line in each plot shows where these place cells (labelled by their row number) have their peak firing location on the full track. Each blue graph represents a particular simulation and shows where each cell has its peak firing location in that simulation. The plots show how the behaviour of the model changes when the track length (*l*), and the strength of grid cell – place cell connections (*G*), are varied.

In the place cell only model (A.12), in contrast, on the two longest of the shortened tracks the shift realignment begins practically straight from the start, even though BVC inputs there are quite weak, and completes in the first half of the journey (Figure 5, *top*). This is in clear contradiction with Gothard et al.’s (1996b) experiment, where on the tracks of similar lengths the realignment only began with the rat approaching the midpoint and completed with it already far into the second half of the journey. On the two shortest tracks the activity bump realigns via jumping also much earlier than jump realignments occured in Gothard et al.’s (1996b) experiment (Figure 5, *bottom*).

**Figure 5.**
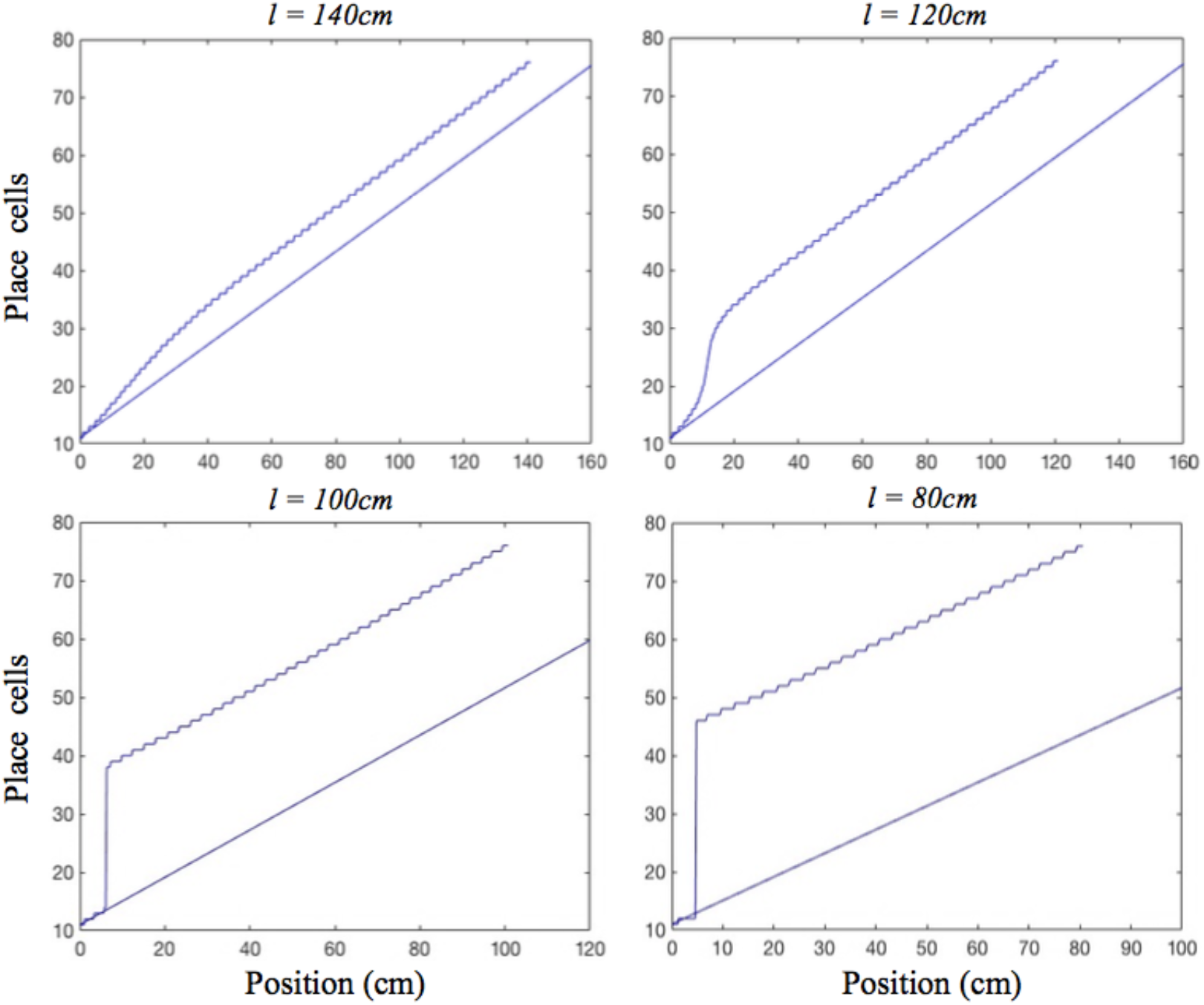
Realignment of the simulated place cell representation of location as track length varies in the place cell only model. Plots show position on the track on the *x* axis and the relevant place cells ordered by their location of peak firing on the full length track (160cm) on the *y* axis. The place cells in rows 11-75 have firing peaks evenly distributed along the full length track from the front of the box. The straight blue line in each plot shows where these place cells (labelled by their row number) have their peak firing location on the full track. Each blue graph represents a particular track length (*l*) simulation and shows where each cell has its peak firing location in that simulation.

We first consider the mechanism behind the dynamics of the place cell only model, and then how the dynamics of the place cell - grid cell model differ from this simpler model. In the place cell only model, the symmetrical connections between place cells (PCs) act to maintain a single activity bump on the PC sheet, whereas asymmetric connectivity, mediated by place by direction cells (Equation A.12), acts to shift the bump, performing path integration (Zhang, 1996; Samsonovich and McNaughton, 1997).

For moderate mismatches (i.e. on the two least shortened tracks) the BVC input to the PC sheet overlaps the activity bump sufficiently to make it shift (with more activity to the leading edge of bump than falling edge), and there are no grid cell projections to stop the bump shifting to align with the BVC input immediately (Figure 5, *top*). In addition, shifts of the PC bump immediately affect the PI input to PCs (as it comes from asymmetric connectivity from the bump itself). For big mismatches (i.e. on the two shortest tracks) the BVC input does not overlap the PC bump, and can cause the bump to jump from the original location to the new location, once the BVC input overcomes the self-support of the initial bump (via the symmetrical and asymmetrical recurrent connections). As the new bump grows it gets the same recurrent support as the original bump, proportional to its size.

For the place cell - grid cell model, the PC activity bump is supported by symmetrical recurrent connections, and recieves inputs from BVCs, but, unlike the place cell only model, the path integrative input to place cells comes from the grid cell (GC) layers, rather than asymmetric connectivity between PCs. In this case, mismatches between BVC inputs and the PC bump have to affect place cell firing before this change can spread to the GC sheets and change their firing before finally affecting the path integrative input to the PC sheet.

For moderate mismatches (i.e. on the two least shortened tracks) when the BVC input overlaps the PC bump sufficiently to make it shift, the PC bump is still receiving a location-specific PI input from the GC layers, preventing any shift faster than that indicated by the rat’s speed. It takes some time for the BVC input to push some of the receiving place cells firing rates above a threshold neccessary for synaptic transmission to the GC layers (see ‘*The place cell – grid cell model’* in the Appendix), which initially produces elongation of the PC bump. When the place cells exceed this firing threshold, they start providing inputs to the GC layers, producing shifting of the GC bumps. Feedback from the additionally shifting GC bumps, finally allows the PC bump to start to shift faster than the rat’s speed to eventually realign with the frame of reference of the approaching end of the track.

The larger the mismatch between the path integration and BVC inputs (i.e. on the second least shortened track versus the least shortened one), the more elongated between the two misaligned inputs the PC activity bump initially becomes, before inducing the realignment of GC bumps by which it is being held back. Therefore, the larger the mismatch, the larger the shifting input to grid cells from place cells, the faster the subsequent activity transition (Figure 4, *top*). During the realignment process, the delay in the feed from the PC to GC layers, the inertia of the GC bumps, and the delay in feedback from the GC to PC layers, all contribute to the pronounced delay before the place cell bump starts shifting, as well as acting as a drag on the speed of its subsequent progress. This produces the shift realignment dynamics as in Gothard et al. (1996b).

For big mismatches (i.e. on the two shortest tracks) the BVC input does not overlap the PC bump and can cause it to jump from its original location to the new one, if the BVC input is sufficiently strong to overcome the self-support of the initial bump and the input from the grid cell modules that holds it in its original location. The BVC input strength, in addition to determining the possibility of realignment, also determines its rapidity, with stronger inputs producing faster place cell activity buildup. Thus, with all values of *G* (2.1, 2.5, 2.8), the bump jumps sooner on the 80cm track than on the 100cm track (Figure 4, *bottom*), since the shorter distance to the end boundary results in stronger BVC inputs. The delay before jump realignments in the place cell – grid cell model (with *G* = 2.1) is much longer than in the place cell only model (as in Gothard et al., 1996b), since, due to the grid cell inputs, greater BVC input buildup is needed for the PC bump to jump. When the bump jumps, it causes the GC bumps to follow, with a slight delay after the place cells realignment. The delay is required for the activities of the grid cells driven by inputs from the realigned PC bump to become strong enough to overcome the initial GC bumps.

In both shift and jump realignments, the dynamics are influenced by the strength of grid cell – place cell connections (*G*). Larger values of *G* increase the influence of grid cell activity on the PC bump, therefore requiring greater BVC input buildup to start the place cell activity transition, leading to a longer initial delay (see Figure 4). The strength of place cell – grid cell connections (*P*) also affects the dynamics of the PC and GC bumps. The value of *P* was set sufficiently high so that changes to the place cell activity affect the GC bumps without significant delay. In shift realignments, this avoids slowing of the rate of shifting of the GC and PC bumps which occurs for low values of *P*. In jump realignments, reducing *P* increases the delay required for the GC bumps to jump, but this delay is relatively short, since, in contrast to the PC bump, only the self-support of the original GC bumps resists their realignment (as for the PC bump in the PC only model jump realignments).

The delays before both shift and jump realignments are significantly longer in the place cell – grid cell model than in the place cell only model. Although the rate of realignment in the place cell only model could be altered by varying parameters such as the relative strength of BVC inputs versus recurrent connectivity, this model cannot show the initial delay before the activity bump starts shifting, as seen in the continuous realignment data. The realignment behavior of the place cell only model is similar to that of the place cell – grid cell model with *G* = 0. In addition, the place cell – grid cell model jumps occurred much earlier on the track than shift realignments (in the middle of the first half of the track versus the second half, see Figure 4 for *G* = 2.1), as seen in the data (Gothard et al., 1996b; Figures 7 and 8). This is because in jump realignments the model place cells initially realign alone, via their activity bump jumping, with grid cell bumps jumping after. In shift realignments, on the contrary, place cells have to continuously pull along resisting grid cell inputs, which slows the realignment. In the place cell only model, both types of realignment occurred on the first half of the track, soon after the start, as the place cells always realign alone. The inability of the place cell only model to show the observed large difference between jump and shift realignment locations provides further support for a parallel place cell – grid cell architecture.

### 4.2 Detailed analysis of Redish et al.’s (2000) data

Redish et al. (2000) investigated which of the following four hypotheses provides the best explanation for the dynamics of hippocampal activity realignment:

(1) Rats use path integration for a certain distance from their starting location, predicting that realignment locations should be distributed as a Gaussian in the box-aligned coordinate frame; (2) Rats use path integration until the room-cues, like the barrier at the end of the track, are percieved, predicting that realignment locations should be distributed as a Gaussian in the room-aligned coordinate frame; (3) Specific landmarks exert control over their own local space, predicting that realignment locations should occur at approximately the halfway point of the journey. (4) Place cells are part of a dynamic system in a semi-stable state that can cross into another semi-stable state only after surpassing an energy barrier, predicting that realignment should at least partially depend on the time elapsed since the mismatch between the sensory and self-motion cues occurred.

For 20 time windows throughout the journey, the first window where place cell activity became more coherent with the room-aligned than box-aligned coordinate frame was determined. This was considered to be the transition point at which the realignment occurred. The consistency of the distribution of the transitions was measured with respect to: (1) the box-aligned coordinate frame; (2) the room-aligned coordinate frame; (3) the midpoint between the box and the barrier; (4) the time since the rat began the journey. The mutual information between each of the four domains and the transition occurrence was measured, to see which of the four would be a more consistent predictor of the realignment occurrence.

The elapsed time since leaving the box was found to be a more consistent predictor than any of the three spatial parameters investigated, suggesting that the shift between coordinate frames is at least partially governed by a time-dependent process. Redish et al. explained the temporal delay before realignment as a stochastic switch occurring somewhere in the system, the stochasticity reflecting noise in the neural network. They concluded that the time preceding the transition reflected the accumulation of sufficient energy for the switch to occur, i.e. for an energy barrier to be surpassed. It was noted that some combination of the four hypotheses could predict the data more completely. The nature of the switch and the specific mechanism and time course of the activity transition itself were left for future research. Below, we address these questions by interpreting the Redish et al. data in terms of the jump and shift mechanisms in our simulations.

Figure 6 shows the distribution of realignment points over all outbound journeys (on different track lengths within the 90-150cm range) of a single animal in the Redish et al. study. Different histograms show distributions of the same realignment points as functions of the four different hypothesized variables. The points of realignment occurrence on the four shortened track lengths from our simulations (with *G* = 2.1) are also shown for comparison in all four different coordinate frames.

**Figure 6.**
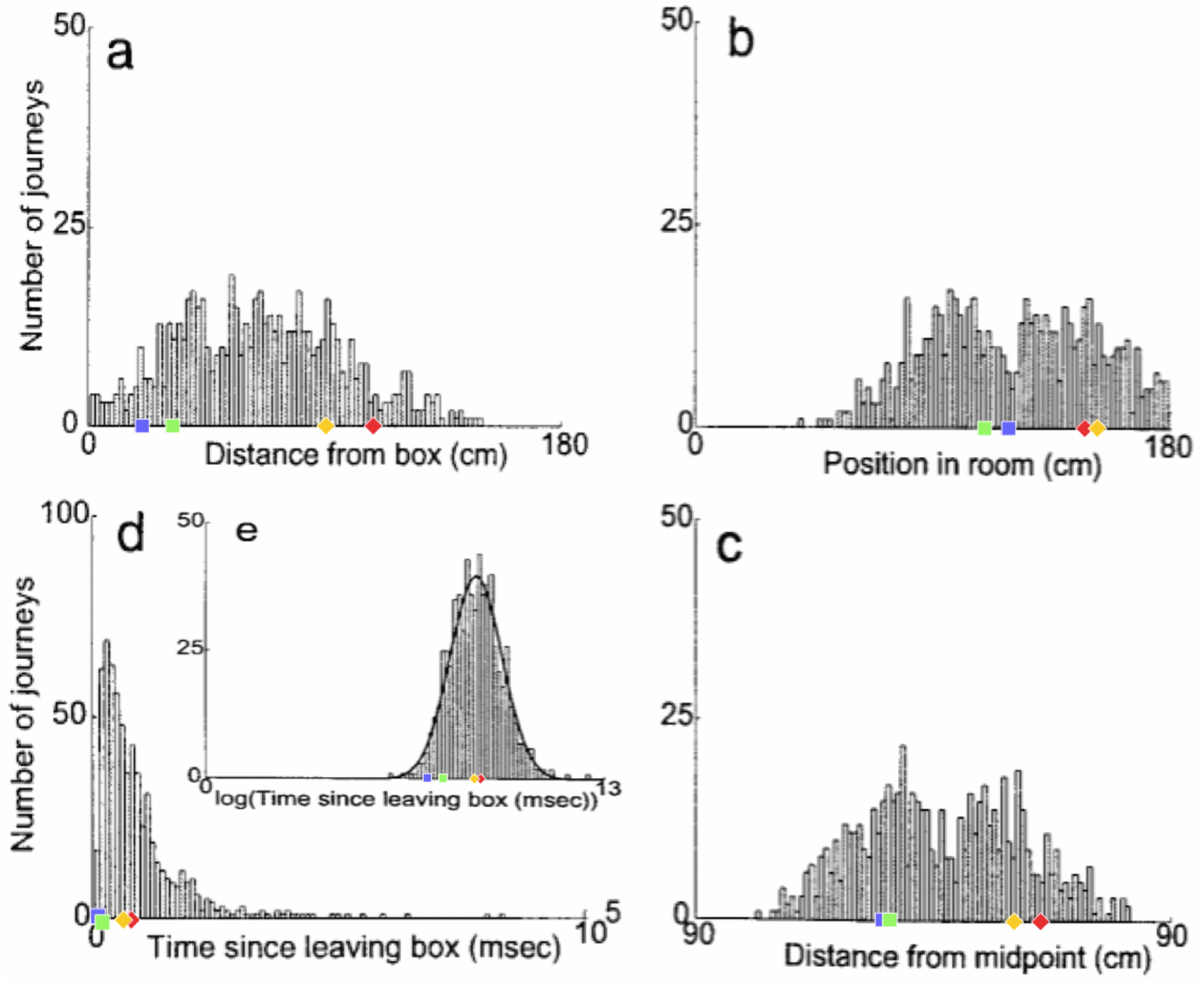
The distribution of realignment points over all outbound journeys from a single animal. a) Distribution of realignments as a function of distance from the box. b) Distribution of realignments as a function of position within the room. c) Distribution of realignments as a function of midpoint between box and barrier. d,e) Distribution of realignments as a function of time since the animal began the outbound journey: d) Time plotted linearly. e) Time plotted logarithmically. A Gaussian function has been fit to the distribution of transition times with time plotted on a logarithmic axis (e, heavy line). Adapted from Redish et al. (2000). Overlayed over the histograms are shown the activity realignment points from our simulations on different track lengths: 140cm in Red; 120cm in Yellow; 100cm in Green; 80cm in Blue. Diamonds indicate continuous (shift) realignments, whereas discontinuous (jump) realignments are represented by squares.

Histogram 6a, with a coordinate system aligned to the movable box, appears to consist of two main overlapping groups of journeys, a narrower group closer to the box (comprising around 40% of journeys), and a much broader group farther from the box. The first group likely comprises jump realignments, since the group spans the range of distances from the box where jumps occur in Gothard et al.’s study and in our simulations. These journeys likely occurred mostly on tracks in the 90-110cm range, since these lengths give the large mismatches that cause jump realignments according to Gothard et al.’s data and our simulations (with *G*=2.1, blue and green dots in Figure 6a). In our simulations, jump realignments occur at approximately 20cm from the box on the 80cm track, and at 32cm on the 100cm track. Thus the observed realignment points forming the peak between 25-40cm (from the box) are consistent with jump realignments in the model, given the differences between the path lengths used and simulated. Earlier realignments (<25cm) likely occurred on the journeys where the rat initially moved very slowly. The second group in Figure 6a consists of journeys where realignment occurred farther away from the box, and thus likely via a shift. The wider realignment points spread in this group is due to a wider range of track lengths with shift realignments (~110-150cm), resulting in a wider range of BVC input strengths experienced across different journeys, and to a range of mismatch distances to be covered by shifting activity on different journeys.

In the time histogram 6d, the mode and adjacent slices likely represent jump realignments, which have shorter delays, and the wider-spread shift realignment points compose the right slope and the tail of histogram 6d. The histogram peaks at shorter delay times because of lower delay time variability in jump than in shift realignments. Slightly greater variability of the distance passed (in 6a) than the delay times (in 6d) before the jump realignment occurrences could be explained by varying speed and trajectory of the rat. Also, at the beginning of longer tracks the strength of BVC input may be insufficient to enable realignment, which, if the rat on these tracks sometimes progressed very slowly in the beginning, could account for the long tail of 6d.

Histogram 6b, with a coordinate system aligned to the end barrier and the room, also shows two distinct groups. The left group should primarily be due to jump realignment cases, since shift realignments mostly occur past the midpoint (which is 75cm from the end for the longest track), according to Gothard et al.’s data and our simulations. The second group, on the contrary, starts around the point where typical jump realignments are expected to stop (see 6b, green and blue dots) and covers the distance range where shift realignments should typically occur (see red and yellow dots in 6b), thus likely representing these. The realignment location variability in both groups is caused, apart from various noise, by the variation in track length across the journeys (e.g. BVC input buildup starts farther from the end on longer tracks).

Consistently with histograms 6a and 6b, histogram 6c is also composed of two salient groups. The sharp right edge of the left group is next to the tracks midpoint, so the group likely consists mostly of jump realignments. Some shift realignments may also contribute to it (e.g. due to rat speed variability), but the bulk of these should compose the right group located after the midpoint. The data shown visibly matches our results (see 6c), as well as Gothard et al.’s data. The presence of both jump and shift realignments is one of the main factors responsible for the wide spread and seeming inconsistency of realignment locations in all three spatial histograms.

Thus the delay in place cell activity realignment, attributed to a stochastic switch by Redish et al. (2000), may result from a parallel place cell – grid cell architecture, as discussed above. Specifically, the variation in the delay across the 90-150cm range of shortened tracks in Redish et al. (2000) is consistent with the delay variation across a similar 80-140cm range of track lengths in our place cell – grid cell model simulations. On the contrary, a single recurrent network of place cells appears to be insufficient to produce similar delay variation across the range of track lengths, in particular the large difference in delays before jump and shift realignments.

Redish et al. (2000) suggested that the variability of the delay prior to realignment reflects noise in the system, using a log-normal distribution to fit the delays across all track lengths (see Figure 6e), although they noted that a hybrid hypothesis may predict the data more fully. Instead, the variation in delay may reflect variations in track lengths and mismatch magnitudes across trials, which are the main predictors of realignment dynamics in our simulations. The further the rat from the end barrier, the weaker the BVC inputs from it, and thus the rate of place cell activity buildup, which leads to longer delays on longer tracks (Figure 6a,d). Also, at the beginning of longer tracks the strength of BVC inputs may be insufficient to override the competing grid cell input to place cells. Additionally, the realignment happens much faster in jump realignments, caused by large mismatches on much shortened tracks, than in continuous (shift) realignments caused by smaller mismatches on moderately shortened tracks (Figure 6a,d).

### 4.3 Individual neural responses during realignment: place cells

The place cell – grid cell model (A.11) can also account for the changes in firing fields of individual place cells on different outbound journeys, as seen in Gothard et al.’s (1996b) study. In the experiment, outbound-selective place cells 2 and 3 showed reduction in size of their firing fields as the track was progressively shortened, and did not fire on the shortest track (Figure 7). The place cells that were active near the box and the track end did not show such changes in firing fields and maintained their position relative to the nearest end of the track across the different track lengths.

**Figure 7.**
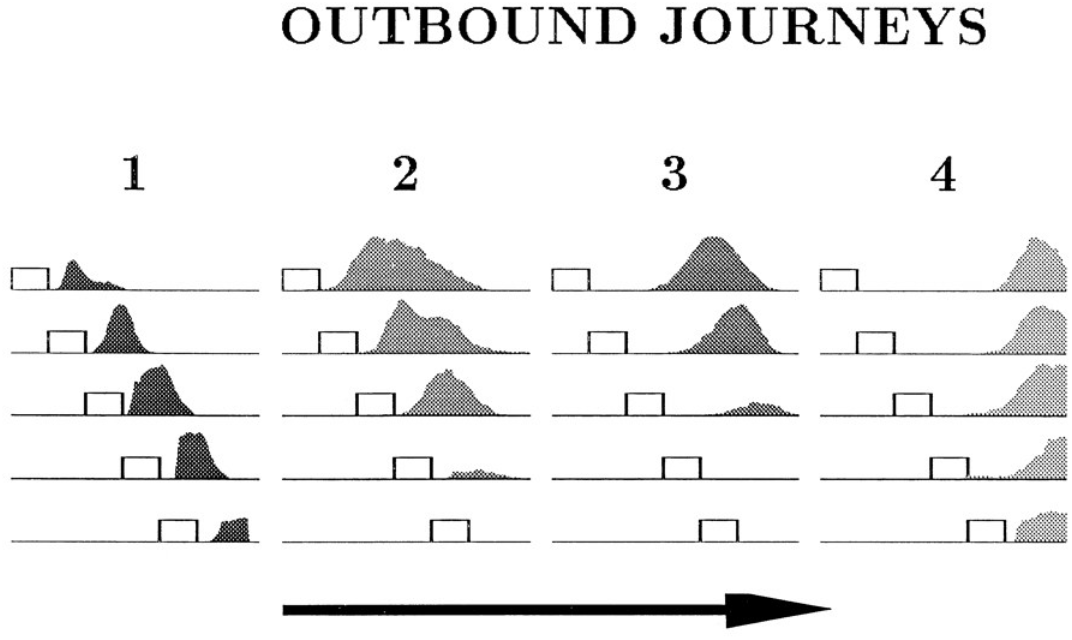
Firing profiles of four outbound-selective cells (1, 2, 3, 4) shown for all five lengths of outbound journey (running direction is indicated by the arrow). The horizontal lines represent the track, and the small rectangles represent the box. The last 27 cm portion of the track, containing the fixed reward cup, is omitted. Cell 1 fired immediately after the rat exited the box, cell 2 fired farther away from the box, cell 3 fired approximately halfway between the box and the end of the track and cell 4 fired close to the end of the track. Adapted from Gothard et al., (1996b).

Figure 8 shows the plots of simulated place cell firing fields on the different track lengths, which qualitatively resemble those of Gothard et al. (1996b). The firing fields centred towards the start and end of the track (Figure 8A, cells 1 and 4) maintain their location relative to the nearest end of the track across the different track lengths. On the shortest track, the firing field near the start of the track (Figure 8A, cell 1) overlaps the location of discontinuous realignment of population activity (see Figure 4) – resulting in curtailment of the right hand side of the firing field. The firing field of Gothard et al’s cell 1 is also reduced on the shortest track compared to longer tracks.

**Figure 8.**
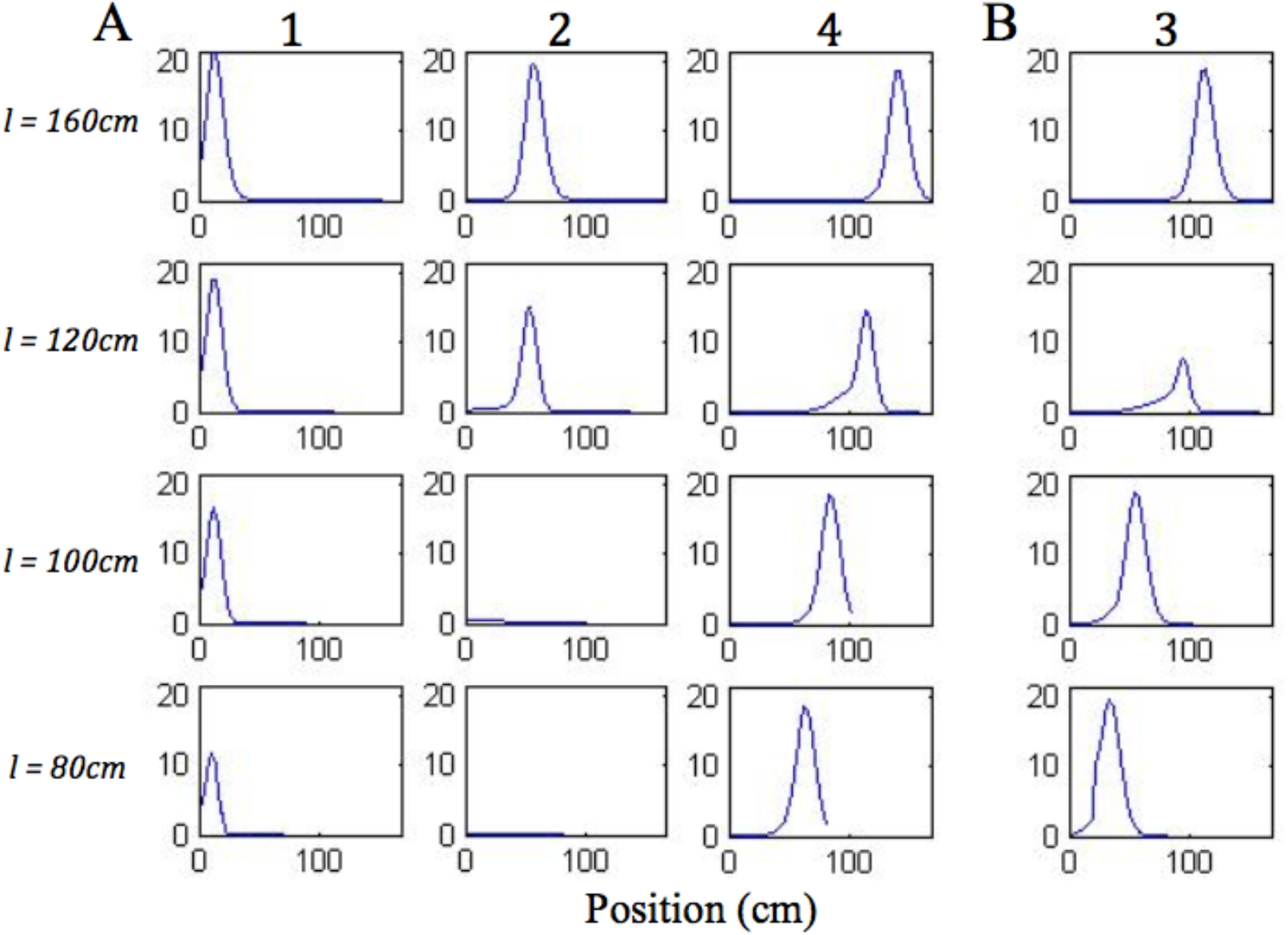
Firing profiles of four illustrative simulated place cells (1, 2, 3, 4) shown for four out of five lengths of outbound journey (the familiar 160cm track, 120cm, 100cm, 80cm). The distance from the box is shown on the x axis in all the plots. Cell 1 fired right after the journey start, cell 4 fired close to the end of the track, and the firing locations of cell 2 and cell 3 are approximately equally spaced between them on the full length track.

The firing field centred at approximately 1/3 of the full length track (Figure 8A, cell 2) shows large changes as the track shrinks. The field of cell 2 narrows on the third shortest track (120cm) and the cell does not fire on the two shortest tracks, since the activity bump jumps over it when realigning there. Gothard et al’s cells 2 and 3 show a similar behaviour.

The field centred at approximately 2/3 of the full length track (Figure 8B, cell 3) largely decreases on the 120cm track, this corresponds to the location of the rapid continuous realignment of population activity (see Figure 4). Our cell 3 firing then recovers on the 100cm track, since the cell fires closer to the end than is the jump realignment location on the track. The cell 3 firing is partially affected by the jump realignment on the shortest track (80cm) – its firing field on the track has a sharp cut off at the left side. The cell 3 behaviour through the progressive track shortening provides a prediction for the pattern of behaviour of the cells that in the Gothard et al.-like experimental conditions on the full length track fire around the firing location of our cell 3 (between Gothard et al.’s cells 3 and 4).

### 4.4 Individual neural responses during realignment: grid cells

What happens to grid cells during the place cell realignment? This experiment (recording grid cells in the situation of Gothard et al., 1996b) has not yet been performed, to our knowledge. However the model presented here makes a clear prediction: because we assume that place cell firing is used to reset the otherwise path-integration driven firing of grid cells. Thus, there should be a smooth compression of the grid in the region of the smooth realignment of place cell firing on the slightly shortened tracks, and a significant disruption or discontinuity in firing of grid cells around the location of the abrupt jump realignment of place cell firing on the shortest tracks.

Figure 9 shows single run simulations of three grid cells, one from each module with different grid scale in the model, on the full length track and four shortened track lengths. On the 140 cm track, a gradual realignment takes place on the second half of the track, with the realignment close to completion by the end of the track. The distance between two neighbouring peaks gets shorter and individual fields get narrower. On the 120 cm track, a rapid shift realignment takes place in the second half. During the rapid realignment, when there is a strong decrease in place cell firing rates, there is a simultaneous increase in corresponding grid cell firing rates. This indicates an increasing input from place cells (due to elongation of the PC activity bump), which results in a more rapid shift of grid cell activity bumps. On the two shortest tracks, grid cell activities jump to a new location soon after the place cell activity jump. As can be seen from the figure, the jumping in all three modules of grid cells happens at the same location, i.e. around the time of the place cell activity jump.

**Figure 9.**
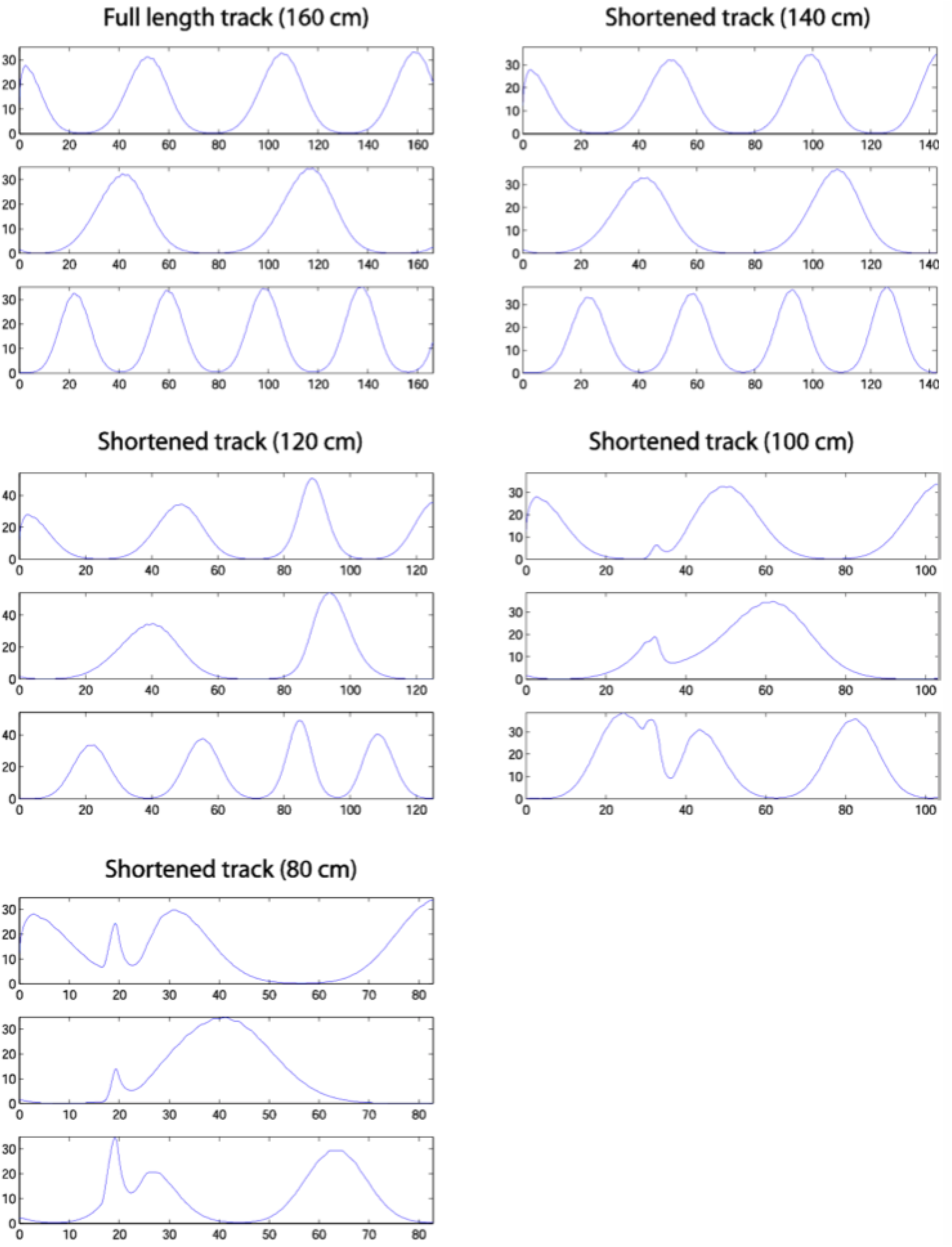
Firing profiles of three simulated grid cells from the three modules with different scales (the ratio between successive grid scales is 0.72). Each plot corresponds to a simulation on the track length shown above the plot, with the same cells shown in the same rows in each plot. Smooth distortion of the grids is seen on the 140cm track. The same effect is seen on the 120cm track, but with considerable changes in peak firing rate also. The two shortest tracks, on which the place cell representation shows discontinuous realignment (see Figure 4), show clear discontinuities in grid cell firing around the location of place cell realignment – indicating a clear effect on the grid cells of the place cell realignment.

These results provide a clear experimental prediction of the model, although the problems of averaging experimental data trial-by-trial, when the location of place cell realignment may vary from trial to trial will need to be borne in mind.

## 5. Discussion

We have presented a detailed description of the mechanisms hypothesised to underline the integration of self-motion and sensory information into a representation of an estimated position. Our navigational model consists of two coupled complementary subsystems working in competition, with the grid cell subsystem specialising in continuous attractor-based path integration and the place cell subsystem in sensory-driven navigation (via Boundary Vector Cells). The model reproduces well the place cell firing rate data from the experiments with conflicting sensory and self-motion inputs, which supports the plausibility of the hypothesised architecture.

Our simulations of the place cell – grid cell model provide explanations for the experimental results of Gothard et al. (1996b) and Redish et al. (2000), and grid cell firing predictions that can be tested in future studies.

### 5.1 The place cell – grid cell system plausibility and potential advantages

The hypothesised model architecture is supported by comparison to the simulations of our place cell only model, which is unable to reproduce Gothard et al. (1996b) and Redish et al. (2000) realignment dynamics.

Sheynikhovich et al. (2009) also simulated Gothard et al.’s (1996b) experiment with conflicting sensory and motion inputs, using their model that, like the place cell only model, does not contain competing semi-autonomous place cell and grid cell subsystems. Instead, modules of recurrently connected grid cells perform path integration as well as receive sensory (visual) inputs directly (like place cells in the place cell only model), and place cells are simply used for read-out. The model shows some of the observed realignment effects (e.g. abrupt activity jumping), but does not fully replicate the Gothard et al. (1996b) and Redish et al. (2000) realignment dynamics. For example, in Sheynikhovich et al.’s model continuous realignments occur before the midpoint even on the longest of shortened tracks. We think that single-layer attracor models cannot show the correct dynamics, as in our place cell only model.

Byrne et al. (2007) also simulated the effects of environment manipulation on place cell firing with their model, which also did not fully reproduce the Gothard et al. (1996b) and Redish et al. (2000) realignment dynamics. As in the place cell only and Sheynikhovich et al. (2009) models, there is one common attractor base, formed by recurrently connected place cells, which receive inputs from BVCs as well as provide location information to a path integration circuit for spatial updating. Thus the path integration circuit does not have its own independent influence on dynamics that would capture the dynamics observed after environmental manipulations.

In contrast to the single-attractor models, the coupled place cell – grid cell system captures the dynamics caused by mis-matching path integration and sensory inputs due to a competitive interaction between two subsystems with semi-independent dynamics.

It is also physiologically plausible for environmental sensory inputs (e.g. provided through boundary related firing: Lever et al., 2009; Solstad et al., 2008) to preferentially control place cell firing, while grid cell firing patterns predominantly reflect self-motion. Recordings in mice running in virtual environments while changing the gain between physical motion and visual motion reveals strong sensory influence on place cell firing patterns compared to much greater influence of self-motion on grid cell firing patterns (Chen et al., 2019). This, together with the evidence for place and grid cells interconnectivity (Brun et al, 2008) and the evidence for excitatory drive from the hippocampus to grid cells (Bonnevie et al., 2013), support the outlined architecture of our model. For grid cells to receive sensory inputs via place cells (rather than directly) also has an advantage of keeping activity patterns in different grid cell modules aligned when visual inputs are not available, otherwise they may quickly lose coherence due to errors accumulating in different modules, leading to place cell activity degradation.

Hardcastle et al. (2015) suggested, based on the results of their experiment, that path integrating grid cells get reset near borders by direct inputs from border cells. The suggested mechanism was recently implemented in the model of Keinath et al. (2018). However, it seems unable to replicate the Gothard et al. (1996b) data as realignment occurs only near the end on all track lengths in their model. In contrast, in our model, grid cells can be reset in any place where adequate boundary-related information is available, via inputs from place cells driven by BVCs, rather than border cells that fire only near borders.

In the recent experiment by Campbell et al. (2018), a mismatch in gain between physical and visual motion was created in mice navigating a virtual linear track (see also Chen et al., 2019). The grid cell behavior was found to depend on the amount of mismatch between self-motion and visual motion. Thus, weaker disagreements caused grid cell firing patterns to shift relative to baseline, as if grid cells constantly tried to align with visual inputs. Larger disagreements, in contrast, led to grid cells breaking free from the influence of landmarks, changing scale relative to them. These findings show a certain similarity to our simulations, where grid cell behavior was also found to depend on the amount of mismatch between self-motion and sensory information, with smaller mismatches resulting in continuous corrective shifting of grid cell activity pattern, while larger mismatches produced discontinuous changes in the activity pattern. In the latter case, our grid cells also for a brief time become free from an influence of environmental sensory inputs, in similarity to Campbell et al. (2018), since the PC activity bump jumps first and therefore the link between the PC and GC activities gets temporarily broken.

The model consisting of two coupled specialist subsystems has certain advantages over models where sensory inputs go directly to a single continuous attractor network that also does path integration, such as place cell-based (e.g. Samsonovich and McNaughton, 1997) or grid cell-based (e.g. Sheynikhovich et al, 2009; Ocko et al., 2018) models. In our model different subsystems serve different purposes and complement each other. The mediation of path integration by recurrent connections between grid cells leaves the recurrent connections between place cells in area CA3 free for other purposes, such as the formation of different continuous attractors for different environments. Thus the place cells can remap between different environments, and perform pattern completion within these remapped representations to accommodate minor sensory changes or cue removal (Wills et al., 2005; Nakazawa et al., 2002) on the basis of the CA3 recurrent connections, independent of the recurrent connections between grid cells. Given that the grid cell representation does not remap, which facilitates development of appropriate recurrent connections for path integration, it is useful to have a separate place cell representation to associate to salient locations. Place cells tend to code for single locations, enabling unambiguous representation of a single goal location, and the place cell representations remap, which allows different goal locations to be represented in different environments.

### 5.2 Mechanism and time-course predictors of hippocampal map realignment

According to Redish et al. (2000), a temporal delay preceding the realignment of place cell population activity in their variable track length experiment could at least partially be explained by a stochastic switch between two semi-stable states. It assumes that place cell population firing patterns form a dynamic system that can only cross into a new state after accumulating enough energy to surpass an energy barrier.

According to our model, the switch effect can be explained by grid cell projections to place cells, which must be overpowered by contradicting sensory inputs in order for the realignment to occur. The sensory stimulus required for inducing the switch is provided by boundaries ahead of the rat via BVC inputs to place cells, which take time to build up sufficiently for the place cells to overcome the path integration influence from grid cells. This causes an initial temporal delay before the population activity transition starts. The complete switch process includes grid cell resetting by place cells (through place cell – grid cell projections), either via grid cell activity packets shifting (concurrently with place cell activity shifting), or via grid cell activity jumping (following place cell activity jumping).

The realignment dynamics are determined by the length of the shortened track on a particular journey. The shortened track length defines the rat’s starting distance to the boundaries ahead (and thus forward-tuned BVC inputs’ strength) and the mismatch magnitude (equal to the amount of track shortening), which determine the initial delay (increasing with the track length) and the time-course of the population activity transition. The larger the mismatch magnitude, as the track length is progressively shortened, the more rapid the continuous activity shifts, which turn into jump transitions when the mismatch magnitude increases above a certain value. Jumps typically occur much earlier on the track than shift realignments, because, according to our model, in jump realignments place cells initially realign alone, with grid cell activities following slightly later. In shift realignments, on the contrary, place cell activity continuously pulls along grid cell activity, which slows the realignment.

### 5.3 Modelling implications and predictions for future research

The results of our simulations show that an architecture of interconnected recurrent networks of place cells (driven by environmental BVC inputs) and grid cells (implementing path integration) can explain the spatial firing patterns and temporal dynamics seen in place cell firing rate data from experiments where the familiar correspondence between environmental and self-motion inputs is manipulated (Gothard et al., 1996b; Redish et al., 2000). In contrast, the alternative architecture of a single recurrent network of place cells (implementing path integration as well as receiving BVC inputs) cannot reproduce these dynamics. Both results support the hypothesis that grid cells and place cells are organised in such a way as to allow two different representations (self-motion and sensory based) to interact with each other. The simulations provide insights into the specific neural mechanisms enabling integration of these two types of information to estimate location.

The simulated place cell – grid cell model makes neuron firing predictions that could be tested in future animal studies. For example, it would be interesting to conduct experiments in rats, similar to those of Gothard et al. and Redish et al., but recording from grid cells as well as place cells. It should be possible to verify that the realignment is driven by a conflict in which sensory input from ahead of the rat eventually resets the path integrative mechanism implemented by grid cells. If so, the model predicts that initially the realignment should start amongst place cells, with the activity realignment dynamics predicted by the track length and amount of shortening (as described above). And the grid cell activity should either realign concurrently with the place cell activity shifting or soon after the abrupt place cell activity jump, depending on the amount of track shortening. The realignment should occur simultaneously across all the grid cell modules with different spatial scales (due to the inputs from the same group of place cells), as well as coherently among the grid cells within each module (due to their recurrent connectivity).

## Acknowledgements

This work was supported by a Wellcome Principal Research Fellowship, the ERC Advanced grant NEUROMEM, the UK Medical Research Council. We thank Alexander Mozeika and Christian Doeller for graphical assistance.

## Author contributions

DL and NB conceptualised the model and the study. DL designed and implemented the computational model, performed simulations and data analysis. DL and NB wrote the manuscript.

## Appendix

## Implementation of the place cell – grid cell model

## Place cell network dynamics

The dynamics of recurrently connected place cells with global feedback inhibition, resulting in a continuous attractor network, can be described by the following equation:

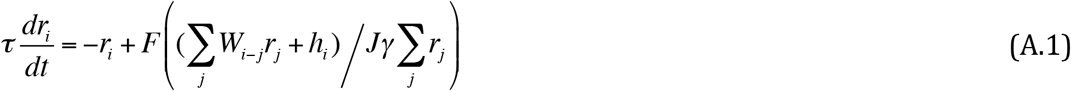

Here *r*_*i*_ is the firing rate of the place cell *i*, *τ* is the place cell time constant, *W*_*i-j*_ is the weight of the recurrent connection from the place cell *j* to the place cell *i*, *h*_*i*_ is an external input and *F* is the response function of the place cells. *γ* represents the strength of the synapse from any place cell onto the global inhibitory neuron and *J* is the synaptic strength of the negative feedback from it (the inhibitory neuron time constant is assumed to be much smaller than that of place cells and is approximated by zero).

We assume the inhibition to be shunting, consistent with the action of shunting GABA synapses (Wilson, 1999). Doiron et al. (2000) have shown that the shunting inhibition has a divisive, rather than subtractive, effect on firing rates of pyramidal cells when they operate at frequencies below 40 Hz. Since place cells fire at lower frequencies, we implement a divisive inhibition in our model. The inhibition is performed through the action of a single, global inhibitory neuron that receives equally strong synapses from all the place cells in the network and provides equal negative feedbacks to all as well, broadly consistent with the very large dendritic and axonal spread of basket cells.

Stringer et al. (2002b) developed a model of Hebbian associative learning of recurrent connections in CA3 during novel environment exploration, which generated a recurrent connections profile with a shape very close to a Gaussian function of the distance between favorite locations of presynaptic and postsynaptic place cells. Therefore we adopt the following function to represent the profile in our model:

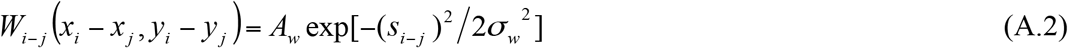

where 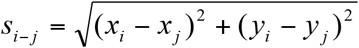 is the distance between the preferred locations of the cells *i*(*x*_*i*_, *y*_*i*_) and *j* (*x*_*j*_, *y*_*j*_), *σ*_*w*_^2^ represents the broadness of the profile and *A*_*w*_ is the maximum weight strength.

We also use the same function to represent recurrent connections between grid cells in the three grid cell modules in our model (A.11), which are also continuous attractor–based. In addition, the function is used to determine the connections between place cells and grid cells in (A.11) (with their profile broadness defined by the average of the variances of the place and grid cell firing fields), since these are assumed to be learnt in a similar Hebbian way.

Since place cell tuning to location often has a shape close to a Gaussian (Hartley et al., 2000), we use the following formula for the place cell tuning curve:

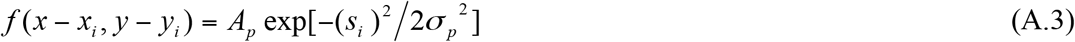

where 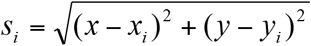 is the distance between the current location of the agent (*x*, *y*) and the preferred location of cell *i*(*x*_*i*_, *y*_*i*_), at which it fires maximally. *σ*_*p*_^2^ is the variance of the place field and *A*_*p*_ is the peak firing rate of the place cell (both assumed to be the same for all place cells in the model for simplicity).

The activation pattern over the population of place cells, virtually arranged so that each one’s location on the Cartesian grid corresponds to its preferred location, is basically equivalent to the place cell tuning curve (Zhang, 1996). We refer to the implicit 2D space represented by the set of activities of so arranged neurons as the ‘state space’ of the system.

If we have a sufficient density of coverage of the represented space by the place cells, in case of a stable activity pattern we can approximate the sum 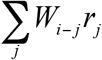 in Equation (A.1) by the 2D-convolution:

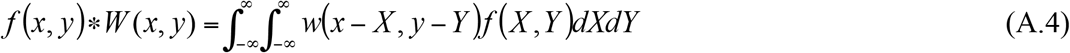

where *f*(*x*, *y*) is the stable firing rate of the place cells given by (A.3).

Since both *W* and *f* are Gaussian, the result of their convolution is another 2D-Gaussian, with the variance *σ*^2^ = *σ*_*w*_^2^ + *σ*_*p*_^2^. In order for Equation (A.1) to be true, our response function *F* has to be a power function of the form:

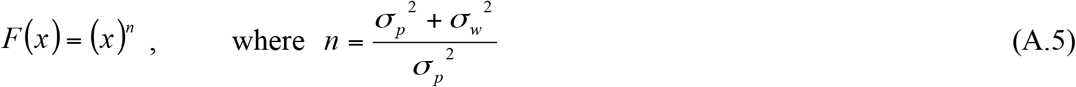

In the case when *σ*_*p*_^2^ = *σ*_*w*_^2^, *F* is simply a square function (n = 2). We can think of this function as a non-saturating region of the Naka-Rushton function, which is considered to provide a reasonably good description of neuronal responses. Thus, with *σ*_*p*_^2^ = *σ*_*w*_^2^, Equation (A.1) becomes:

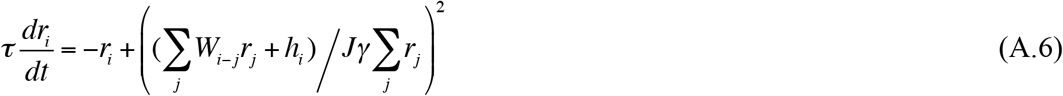

This equation is used as a basis for the description of place cell network dynamics, as well as grid cell network dynamics, since both cell types are assumed to form continuous attractor networks.

## Boundary Vector Cell (BVC) inputs to place cells

In our model every place cell is assumed to receive inputs from four orthogonally tuned BVCs, each of which has directional tuning perpendicular to one of the four surrounding boundaries. The position of the bump-shaped population activity pattern in the state space, representing the environment, is thus controlled by four sets of BVC inputs, each associated with a boundary in a specific direction. Each set of these inputs, or an “input profile”, is composed of the individual inputs from all the BVCs tuned to that boundary.

In our modelling we assume these input profiles to have a Gaussian crossection in the plane orthogonal to their associated boundary, with the mean of the Gaussian occurring at the current distance of the rat from that boundary. We use a Gaussian shape for the input profiles in line with the Gaussian shape of the place cell activity bump, and the point where the peaks of all four profiles overlap corresponds to the activity bump centre. We assume that the amplitude and variance of these Gaussian profiles vary linearly with the distance from their associated boundaries, with the amplitude decreasing and variance increasing, corresponding to the decreasing influence of more distant boundaries.

We then translate a set, consisting of all the input profiles associated with a particular boundary and distributed through 160cm (the original track length in Gothard et al. (1996b)) starting from the boundary, into a set of tuning curves for individual BVCs distributed through that distance. Equation (A.7) has been found to provide the description for the firing response *b*_*i*_ of each individual BVC *i* with its preferred distance *d*_*i*_ as a function of distance to the boundary *x*:

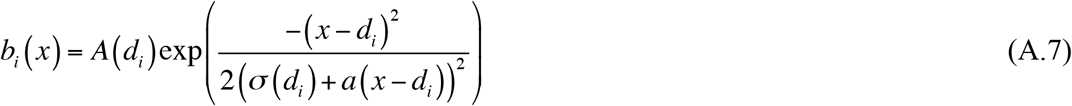

Both *A* and *σ* here vary linearly with variation of the preferred distance across BVCs, i.e. *σ*(*d*_*i*_) = *σ*_0_ + *γd*_*i*_ and *A*(*d*_*i*_) = *A*_*0*_ − *βd*_*i*_, where *γ* and *β* are constants determining respectively rates of *σ* increase and *A* decrease with the preferred distance increase. *α* gives the rate with which *σ* of a particular BVC varies linearly with the difference between its preferred and current distance from the boundary.

On the contrary, if we assumed a Gaussian shape for the tuning curves of individual BVCs, rather than for the overall input profile from a boundary, the resulting overall input profiles would not be Gaussian but skewed, and consequently their combined input (i.e. their sum) to the place cell activity bump would not be well focused. Although in the case of single place cells a Gaussian BVC curve may give a reasonable fit, as in O’Keefe and Burgess (1996), it does not work so well for a whole group of place cells.

The total external input *h*_*i*_ to the place cell *i* we model as proportional to the thresholded linear sum of all BVC inputs *b*_*j*_ (see (A.7)) it receives:

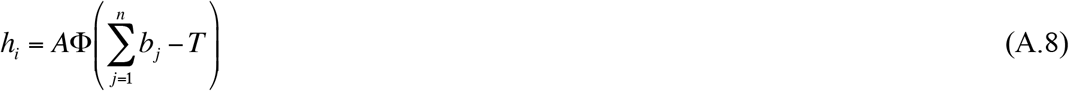

where the weighting parameter *A* is constant, *T* is a threshold, and Φ is the Heaviside function (i.e. Φ(*x*) = *x* if *x* > 0, Φ(*x*) = 0 otherwise).

## Conjunctive grid by head-direction cell responses

In order for each grid cell to have asymmetric connections to the six grid cells with firing patterns offset in 6 directions, mediated by conjunctive cells with the corresponding directional preferences, there need to be six distinct groups of grid by head-direction cells, each composed of cells tuned to one of the six directions. The number of cells in each group is equal to the total number of grid cells, so that each grid cell has a corresponding (i.e. with the same grid node locations) grid by head-direction cell in each of the six groups. We define the firing response *g*_*jD*_ of the grid by head-direction cell *j*_*D*_ from the group *D*, consisting of cells with the preferred head direction *θ*^*D*^, as follows:

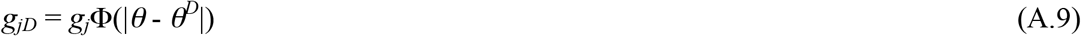

where *g*_*j*_ is the firing response of the corresponding grid cell *j* (which may potentially be providing a feed-forward input to the grid by head-direction cell *j*_*D*_). The function Φ(of rat’s head direction *θ*) describes the directional modulation of the grid by head-direction cell responses in the following way:

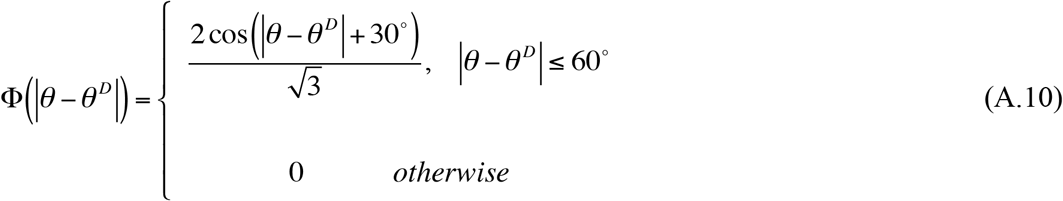

Equation (A.10) provides the form of modulation of conjunctive cell inputs to a grid cell that is necessary for correct path integration, i.e. so that the sum of different translation vectors (produced by different conjunctive cell inputs to a grid cell) was always equal to the actual rat’s translation. (In the case of 6 directions, conjunctive cell firing modulation by (A.10) has the same overall effect as the modulation by a cosine function has in a Cartesian coordinate system (4 directions).)

## The place cell – grid cell model

The proposed place cell – grid cell system is represented by the following set of equations (based on Equation (A.6)):

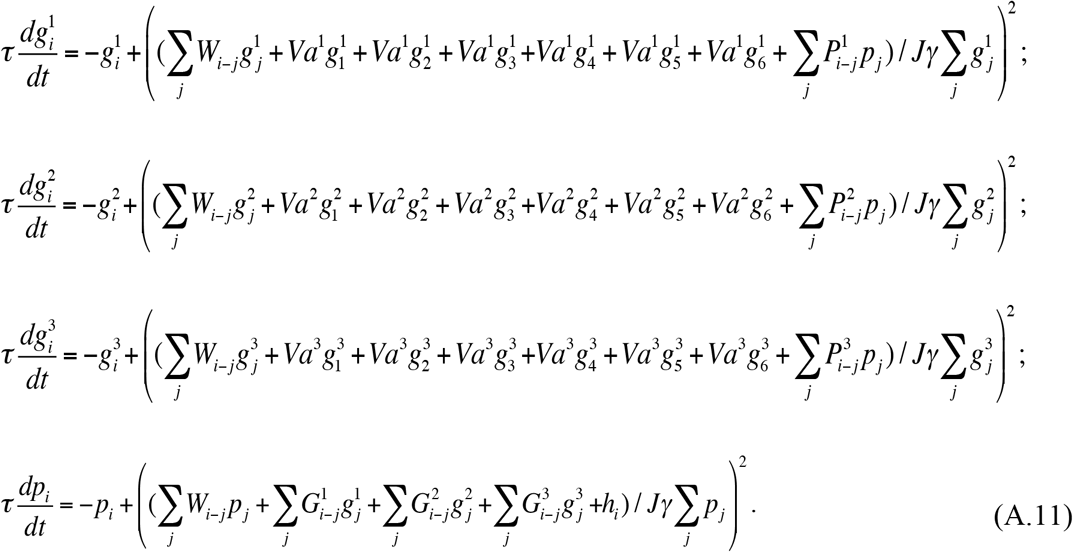

Here 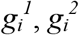 and 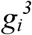 are the firing rates of the grid cells *i* from the first, second and third module of cells correspondingly, whereas *p*_*i*_ is the firing rate of the place cell *i*. *W*_*i-j*_ is the weight of the recurrent connection from the grid/place cell *j* to the grid/place cell *i*, given by (A.2). *P*_*i-j*_ is the connection weight from the place cell *j* to the grid cell *i* and *G*_*i-j*_ is the connection weight from the grid cell *j* to the place cell *i*, both determined according to (A.2). 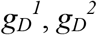 and 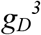 are the firing rates of the grid by head-direction cells from the first, second and third module correspondingly, given by (A.9) with *D* = 1,2,3,4,5,6 and *θ*^*D*^ = 0°,60°,120°,180°,240°,300°, which corresponding grid cells are located (in the state space) next to and in the direction *θ*^*D*^+180° from the grid cells *i* (in the three modules). *a*^*1*^, *a*^*2*^, *a*^*3*^ are the weights of the input to the grid cell *i* from each of the six grid by head-direction cells in the first, second and third module correspondingly. *γ* represents the strength of a synapse from any grid/place cell onto the global inhibitory neuron for its module of grid/place cells, and *J* is the synaptic strength of the negative feedback from it. *V* is the speed of rat’s movement. *h*_*i*_ is the external input to the place cell *i*, given by (A.8). The synaptic strengths *γ* and *J* are respectively set to 0.04 and 0.123 for each grid cell module and place cells, and the model time constant *τ* is set to 0.05.

An assumption was made regarding synaptic transmission between the place and grid cell layers: that only high firing rates propagated through the connections between them. The reason for this was to ensure the input between the layers was spatially specific – i.e. only firing from the centres of the firing fields, and thus activity bumps, of place and grid cells should influence the representation in the other layer. There is some physiological evidence to support the idea that only spikes fired during periods of high-firing rate (“bursts” of spikes) are reliably transmitted by synapses in the hippocampal region (see Lisman, 1997, for a review). Thus, only firing rates exceeding a threshold *r* propagated between the place and grid cells, with *r* = *r*_*max*_ − 6 for the grid cell to place cell projections, where *r*_*max*_ is the highest grid cell firing rate at a particular time. The firing rate threshold for the place cell to grid cell projections was set a bit higher, to: *r* = *r*_*max*_ − 4. We had to raise it higher than for the grid cell projections in order to prevent two largely mismatched inputs (one from the place cell activity bump and the other from the building up sensory input) going to the grid cells simultaneously, which otherwise causes activity bumps oscillation prior to the realignment occurrence on the two shortest tracks, corrupting firing fields located before the realignment point.

## Implementation of the place cell only model

The following Equation (A.12) describes the place cell only model for spatial navigation. The model does not have grid cells, instead path integration is performed by recurrently connected place cells with place by direction neurons:

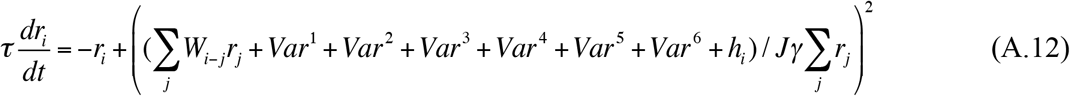

Here *r*_*i*_ is the firing rate of the place cell *i*. *r*^*D*^ = *r*Φ(|*θ* - *θ*^*D*^|), with *D* = 1,2,3,4,5,6 and *θ*^*D*^ = 0°,60°,120°,180°,240°,300° correspondingly, is the firing rate of the place by direction neuron from the group *D* (with Φ given by A.10), which corresponding place cell (with firing rate *r*) is located (in the state space) next to and in the direction *θ*^*D*^ + 180° from the place cell *i*. *W*_*i-j*_ is the weight of the recurrent connection from the place cell *j* to the place cell *i* given by (A.2) and *a* is the weight of the input to the place cell *i* from each of the six place by direction neurons. *γ* represents the strength of a synapse from any place cell onto the global inhibitory neuron for place cells, and *J* is the synaptic strength of the negative feedback from it. *V* is the speed and *θ* is the direction of rat’s movement. *h*_*i*_ is the external input to the place cell *i* given by (A.8) and (A.7) as for the place cell – grid cell model. The synaptic strengths *γ* and *J* are respectively set to 0.04 and 0.123, and the model time constant *τ* is set to 0.05, all as in the other model.

## References

Amaral DG, Witter MP (1989) The three-dimensional organization of the hippocampal formation: a review of anatomical data. Neuroscience 31: 571–591.

Barry C, Hayman R, Burgess N, Jeffery KJ (2007). Experience-dependent rescaling of entorhinal grids. Nature Neurosci 10: 682–684.

Barry C, Lever C, Hayman R, Hartley T, Burton S, O’Keefe J, Jeffery KJ, Burgess N (2006) The boundary vector cell model of place cell firing and spatial memory. Reviews in the Neurosciences 17(1-2): 71–79.

Bonnevie T, Dunn B, Fyhn M, Hafting T, Derdikman D, Kubie JL, Roudi Y, Moser EI, Moser MB (2013). Grid cells require excitatory drive from the hippocampus. Nature Neurosci 16(3): 309–17.

Brun VH1, Leutgeb S, Wu HQ, Schwarcz R, Witter MP, Moser EI, Moser MB. (2008) Impaired spatial representation in CA1 after lesion of direct input from entorhinal cortex. Neuron 57(2): 290–302.

Yoram Burak, Ila R Fiete (2009) Accurate Path Integration in Continuous Attractor Network Models of Grid Cells. PLoS Comput Biol 5(2): e1000291. doi:10.1371/journal.pcbi.1000291.

Byrne P, Becker S, Burgess N (2007) Remembering the past and imagining the future: a neural model of spatial memory and imagery. Psychol Rev 114: 340–375.

Malcolm G. Campbell, Samuel A. Ocko, Caitlin S. Mallory, Isabel I. C. Low, Surya Ganguli, and Lisa M. Giocomo (2018) Principles governing the integration of landmark and self-motion cues in entorhinal cortical codes for navigation. Nature Neurosci 21: 1096–1106.

Conklin J, Eliasmith C (2005) A controlled attractor network model of path integration in the rat. J Comput Neurosci. 18(2):183–203.

Guifen Chen, Yi Lu, John A King, Francesca Cacucci, Neil Burgess (2019) Differential influences of environment and self-motion on place and grid cell firing. Nature Communications 10: 630 (2019).

Dhillon A, Jones RS (2000) Laminar differences in recurrent excitatory transmission in the rat entorhinal cortex in vitro. Neuroscience 99: 413–422.

Doiron B, Longtin A, Berman N, Maler L (2000) Subtractive and divisive inhibition: Effect of voltage-dependent inhibitory conductances and noise. Neural Computation 12:1–22.

Etienne AS, Maurer R, Seguinot V (1996) Path integration in mammals and its interaction with visual landmarks. J Exp Biol 199: 201–209.

Ila R. Fiete, Yoram Burak and Ted Brookings (2008) What Grid Cells Convey about Rat Location. Journal of Neuroscience 28 (27) 6858–6871.

Germroth P, Schwerdtfeger WK, Buhl EH (1991) Ultrastructure and aspects of functional organization of pyramidal and nonpyramidal entorhinal projection neurons contributing to the perforant path. J Comp Neurol 305: 215–-231.

Gothard KM, Skaggs WE, McNaughton BL (1996b) Dynamics of mismatch correction in the hippocampal ensemble code for space: interaction between path integration and environmental cues. J Neurosci 16: 8027–40.

Roddy M Grieves, Éléonore Duvelle & Paul A Dudchenko (2018): A boundary vector cell model of place field repetition, Spatial Cognition & Computation, DOI: 10.1080/13875868.2018.1437621.

Alexis Guanella, Daniel Kiper, Paul F.M.J. Verschure (2007) A model of grid cells based on a twisted torus topology. International Journal of Neural Systems 17(4): 231–40.

Hafting T, Fyhn M, Molden S, Moser M-B, Moser EI (2005) Microstructure of a spatial map in the entorhinal cortex. Nature 436: 801–806.

Hardcastle K, Ganguli S, Giocomo LM (2015). Environmental boundaries as an error correction mechanism for grid cells. Neuron. 2015 May 6; 86(3):827–39.

Hartley T, Burgess N, Lever C, Cacucci F, O’Keefe J (2000) Modeling place fields in terms of the cortical inputs to the hippocampus. Hippocampus 10:369–379.

Alexandra T Keinath, Russell A Epstein, Vijay Balasubramanian (2018) Environmental deformations dynamically shift the spatial metric. eLife 2018;7:e38169.

Julija Krupic, Marius Bauza, Stephen Burton, Caswell Barry & John O’Keefe (2015) Grid cell symmetry is shaped by environmental geometry. Nature 518: 232–235.

Julija Krupic, Marius Bauza, Stephen Burton, John O’Keefe (2018) Local transformations of the hippocampal cognitive map. Science 359, 6380: 1143–1146.

Colin Lever, Stephen Burton, Ali Jeewajee, John O’Keefe, and Neil Burgess (2009) Boundary Vector Cells in the Subiculum of the Hippocampal Formation. The Journal of Neuroscience 29(31): 9771–7.

Lingenhohl K, Finch DM (1991) Morphological characterization of rat entorhinal neurons in vivo: soma-dendritic structure and axonal domains. Exp Brain Res 84: 57–-74.

Lisman JE (1997) Bursts as a unit of neural information: making unreliable synapses reliable. Trends. Neurosci. 20:38–43.

McNaughton BL, Battaglia FP, Jensen O, Moser EI, Moser MB (2006). Path integration and the neural basis of the ‘cognitive map’. Nat.Rev.Neurosci 7: 663–678.

Nakazawa K, Quirk MC, Chitwood RA, Watanabe M, Yeckel MF, Sun LD, Kato A, Carr CA, Johnston D, Wilson MA, Tonegawa S (2002) Requirement for hippocampal CA3 NMDA receptors in associative memory recall. Science 297: 211–8.

Zaneta Navratilova, Lisa M Giocomo, Jean-Marc Fellous, Michael E Hasselmo and Bruce L McNaughton (2011). Phase Precession and Variable Spatial Scaling in a Periodic Attractor Map Model of Medial Entorhinal Grid Cells with Realistic After-Spike Dynamics, Hippocampus 22(4):772–89.

O’Keefe J (1976) Place units in the hippocampus of the freely moving rat. Exp Neurol 51:78–109.

O’Keefe (2007) Hippocampal Neurophysiology in the Behaving Animal. The Hippocampus Book (Chapter 11). Editors: Per Andersen, Richard Morris, David Amaral, Tim Bliss and John O’Keefe. Oxford University Press.

O’Keefe J, Burgess N (1996) Geometric determinants of the place fields of hippocampal neurons. Nature 381:425–428.

O’Keefe J, Burgess N (2005) Dual phase and rate coding in hippocampal place cells: theoretical significance and relationship to entorhinal grid cells. Hippocampus 7: 853–66.

O’Keefe J, Dostrovsky J (1971) The hippocampus as a spatial map. Preliminary evidence from unit activity in the freely-moving rat. Brain Res 34:171–175.

O’Keefe J, Nadel L (1978) The hippocampus as a cognitive map. Oxford: Oxford University Press.

Samuel A. Ocko, Kiah Hardcastle, Lisa M. Giocomo, and Surya Ganguli (2018) Emergent elasticity in the neural code for space. PNAS 115 (50) E11798–E11806.

Redish AD, Rosenzweig ES, Bohanick JD, McNaughton BL, Barnes CA (2000) Dynamics of hippocampal ensemble activity realignment: time versus space. Journal of Neuroscience 20:9298–9309.

Redish AD, Touretzky DS (1998) “The Role of the Hippocampus in Solving the Morris Water Maze”. Neural Computation 10(1): 73–112.

Samsonovich A, McNaughton BL (1997) Path integration and cognitive mapping in a continuous attractor neural network model. J Neurosci 17: 5900–20.

Sheynikhovich, D., Chavarriaga, R., Strösslin, T., Arleo, A., and Gerstner, W. (2009). Is there a geometric module for spatial orientation? Insights from a rodent navigation model. Psychol. Rev. 116: 540–66.

T. Solstad, C. N. Boccara, E. Kropff, M. B. Moser, E. I. Moser (2008) Representation of Geometric Borders in the Entorhinal Cortex. Science 322: 1865–8.

Hanne Stensola, Tor Stensola, Trygve Solstad, Kristian Frøland, May-Britt Moser & Edvard I. Moser (2012) The entorhinal grid map is discretized. Nature 492: 72–78.

Tor Stensola, Hanne Stensola, May-Britt Moser, Edvard I. Moser (2015) Shearing-induced asymmetry in entorhinal grid cells. Nature 518: 207–12.

Stringer SM, Rolls ET, Trappenberg TP, de Araujo IET (2002b) Self-organizing continuous attractor networks and path integration: two-dimensional models of place cells. Network: Computation in Neural Systems 13: 429–446.

Tom J. Wills, Colin Lever, Francesca Cacucci, Neil Burgess, John O’Keefe (2005) Attractor Dynamics in the Hippocampal Representation of the Local Environment. Science 308(5723): 873–6.

Wilson HR (1999) Spikes, Decisions and Actions. The dynamical foundations of neuroscience. Oxford University Press, Oxford. p 126.

Zhang K (1996) Representation of spatial orientation by the intrinsic dynamics of the head-direction cell ensemble: a theory. J Neurosci 16: 2112–26.

